# Bulk and Mosaic Deletions of *Egfr* Reveal Regionally Defined Gliogenesis in the Developing Mouse Forebrain

**DOI:** 10.1101/2022.12.10.519926

**Authors:** Xuying Zhang, Guanxi Xiao, Caroline Johnson, Yuheng Cai, Zachary K. Horowitz, Christine Mennicke, Robert Coffey, Mansoor Haider, David Threadgill, Rebecca Eliscu, Michael C. Oldham, Alon Greenbaum, H. Troy Ghashghaei

## Abstract

The Epidermal growth factor receptor (EGFR) plays a role in cell proliferation and differentiation during healthy development and tumor growth, however its requirement for brain development remains unclear. Here we used a conditional mouse allele for *Egfr* to examine its contributions to perinatal forebrain development at the tissue level. Subtractive bulk ventral and dorsal forebrain deletions of *Egfr* uncovered significant and permanent decreases in oligodendrogenesis and myelination in the cortex and corpus callosum. Additionally, an increase in astrogenesis or reactive astrocytes in effected regions was evident in response to cortical scarring. Sparse deletion using Mosaic Analysis with Double Markers (MADM) surprisingly revealed a regional requirement for EGFR in rostrodorsal, but not ventrocaudal glial lineages including both astrocytes and oligodendrocytes. The EGFR-independent ventral glial progenitors may compensate for the missing EGFR-dependent dorsal glia in the bulk *Egfr*-deleted forebrain, potentially exposing a regenerative population of gliogenic progenitors in the mouse forebrain.

## Introduction

A fundamental question in developmental neuroscience is how the sequential production of diverse populations of neurons and glia in the central nervous system (CNS) is regulated in space and time. The balanced production of both neurons and glia is required for homeostatic functioning of the CNS. Defects in this balance can result in developmental disabilities and brain malformations with severe consequences in newborns and later in life ^**1–3**^. However, mechanisms that regulate the developmental transition of neurogenesis to gliogenesis and its potential spatio-temporal and cellular diversity remain partially understood ^**4**^.

Neurogenesis in the cerebral cortex is characterized by an initial expansion of neuroepithelial cells followed by their transition into neurogenic progenitors ^**5**^. Here, excitatory pyramidal neurons are generated from neural progenitors in the dorsal telencephalic (dorsal pallial) ventricular and subventricular zones (VZ/SVZ) which then migrate basally into the developing cortical plate and form layers in a birthdate-dependent manner ^**6**^. Inhibitory interneurons are generated in the ventral forebrain (or ventral pallium) and migrate tangentially into the cortex during embryonic development ^**7**^. When cortical neurogenesis dwindles in the late-stage embryo, neurogenic progenitors transform into glial progenitors or directly into glia, forming a plethora of critical support systems for neuronal survival and function ^**3,8,9**^. Several waves of proliferation through and after the neurogenesis to gliogenesis switch during late embryonic and early postnatal stages give rise to the majority of glia in the mature cortex ^**10,11**^. Unlike neurogenesis which largely stops during late embryonic development and lacks regenerative capacity, glial expansion continues postnatally, and is responsive to mechanical, ischemic, aging, and disease related injuries in the CNS ^**3,9,12**^.

Heterogeneity among mature glia is evident in the diversity of their developmental origins, morphology, gene expression and function. Two general classes constitute glia in the CNS, macroglia and microglia. Macroglia developmentally arise from progenitors within the CNS (neuroectodermal origin), whereas microglia are derived from the hematopoietic system (mesodermal origin) around midgestation when they enter and seed the CNS from the circulating blood, and function as neuroimmune cells throughout life ^**13**^. Macroglia (astrocytes and oligodendrocytes) are distributed throughout the white and grey matters of the mature CNS where they exhibit regional and molecular diversity ^**14–16**^ and function as supporting cells for various neural networks. Macroglia are derived from 10-20% of neurogenic progenitors in the cortex ^**17,18**^ with astrocytes maintaining their ventral-dorsal identities both in the forebrain ^**19**^ and the developing spinal cord ^**20**^. Development of cortical oligodendrocytes is more complex in that they originate from both the ventral and dorsal telencephalic regions ^**10**^. Genetic programs and molecular markers that distinguish glial cell types have been identified, however much remains to be understood regarding their regional heterogeneity, lineage overlap, and interactions among factors underlying their development and function in the CNS.

Among factors with established roles in gliogenesis are the receptor tyrosine kinases (such as ERBB receptors) and their downstream signals ^**21**^. The first ERBB member is the epidermal growth factor receptor (EGFR; i.e. ERBB1), which is known to be expressed in the periventricular regions of the developing brain ^**22–25**^. While the developmental function of EGFR has been studied in various contexts since the first report on its germline deletion ^**26**^, its precise role in the development of the CNS remains unclear. Gliogenesis is certainly under the influence of EGFR in a dose-dependent manner presumably through its activation by its predominant ligands EGF and TGF ^**24,27**^. A number of studies have implicated EGFR signaling in oligodendrocyte development ^**25,28–31**^, and in embryonic and adult neurogenesis and neuronal survival ^**27,32,33**^. We and others have suggested a role for EGFR in both astrocyte and oligodendrocyte production ^**18,33–36**^, and asymmetric inheritance of EGFR following mitosis in progenitors was shown to demarcate gliogenic and neurogenic lineages ^**37**^. Another study found minimal defects in CNS homeostasis in brain-concentrated (*Nestin-cre*) conditional *Egfr-null* mice, but that defective astrocytes in these mice made them susceptible to Kainate-induced epilepsy ^**38**^. As such, the extent to which EGFR regulates gliogenesis and neurogenesis in the forebrain remains unclear. Here we used region-specific conditional deletions of *Egfr* to characterize its effects on forebrain development in mice. Additional investigation using Mosaic Analysis with Double Markers (MADM) reveals an unexpected gradient of EGFR-requirement for gliogenesis along the rostrocaudal and ventrodorsal axes, indicating the existence of EGFR-dependent and EGFR-independent gliogenic progenitor domains in the forebrain.

## Results

### Time-, region-, and cell-specific expression of EGFR in the developing cortex

We examined the developmental timing of EGFR expression in the mouse forebrain using in situ hybridization panels from the Allen Brain Atlas (ABA), antibody staining of wild type (*WT*) forebrain sections, and mice in which EGFR is expressed with a fluorescent epitope tag, Emerald GFP fusion ^39^ (*Egfr ^EM^*) (**Fig. 1A**). EGFR was primarily expressed in the ventricular zone of the dorsal telencephalon and ventrally in the ganglionic eminences (GE), around embryonic day 15.5 (E15.5). Its expression appeared upregulated between E18.5 and postnatal day 4 (P4) in the dorsal telencephalon (subependymal zone, SEZ; white matter, WM; cortex, Ctx), in the ventral telencephalon (striatum, Str; rostral migratory stream, RMS), and the olfactory bulbs (OB). From P4 onward, EGFR expression became restricted to the SEZ, RMS, OB, and WM (**Fig. 1A**). A thorough examination of the potential EGFR ligands in the forebrain using ABA panels indicated that TGFα, NRG1, and NRG2 exhibit spatial and temporal expression patterns suggesting that they may function as ligands for EGFR during perinatal stages (**Fig. S1**).

**Figure 1.**
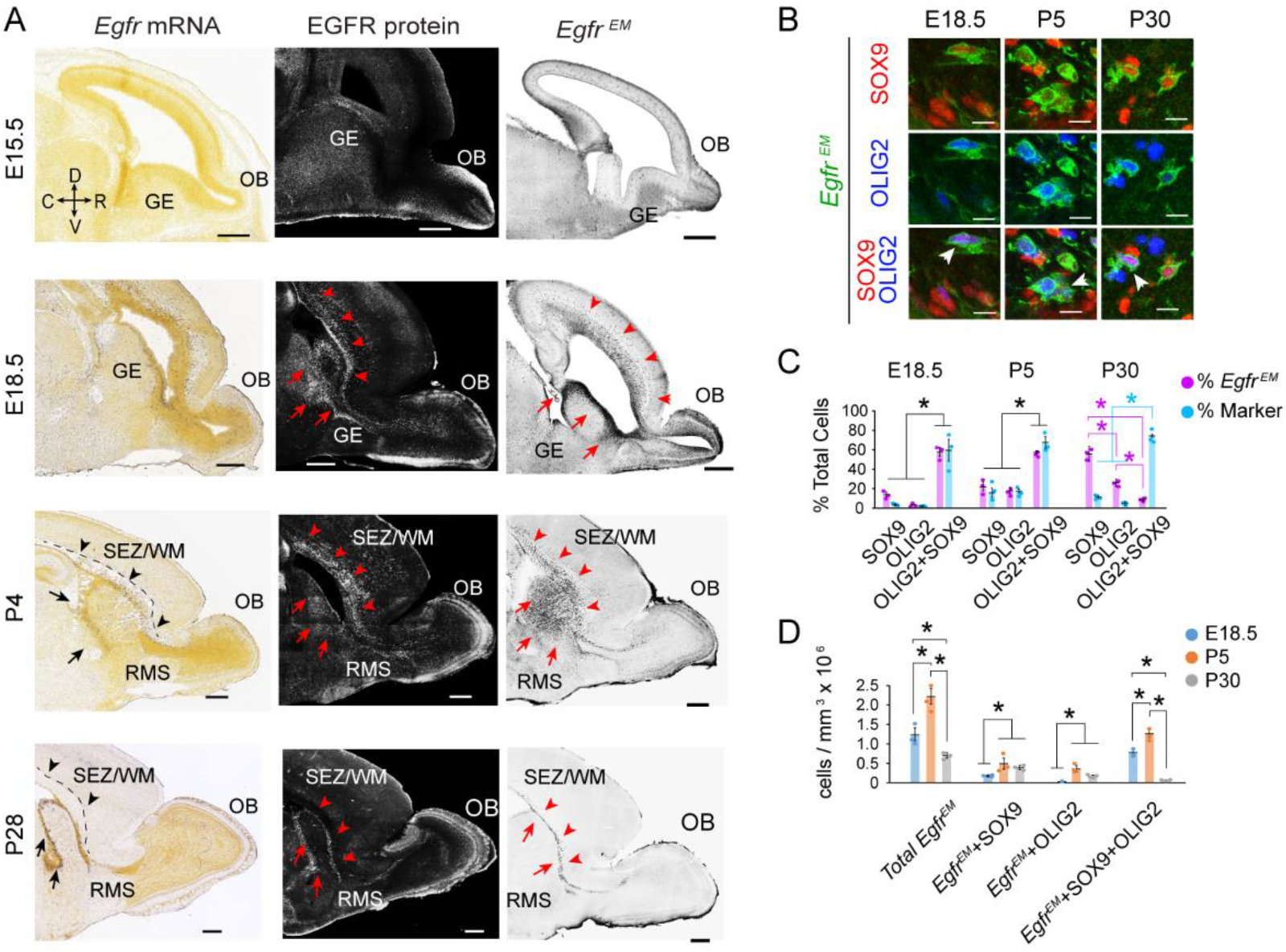
Developmental expression of EGFR in the forebrain. **(A)** In-situ hybridization panels from the Allen Brain Atlas (Developing Mouse Brain) illustrating localization of *Egfr* mRNA in the embryonic and postnatal forebrain (left column). Confocal images of embryonic and postnatal forebrain tissues immunostained for EGFR in *WT* mice (middle column) and in *Egfr^EM^* mice (expressing EGFR fused to the fluorescent protein Emerald GFP, EM; right column). Arrowheads point to the dorsal subependymal zone and white matter (SEZ/WM) and arrows to the ventral SEZ. RMS, rostral migratory stream; OB, olfactory bulb; GE, ganglionic eminence. Ages of mice are as indicated for each row. All images are sagittal views. Scale bars, 500 μm. **(B)** Overlap of Olig2 and Sox9 with *Egfr ^EM^* positive cells in the SEZ/WM. Arrows point to triple labeled cells. Scale bars, 10 μm. **(C)**Analysis of *Egfr^EM^* cells coexpressing SOX9 and/or OLIG2 using percentage of *Egfr ^EM^* cells expressing each or both markers (% *Egfr^EM^*), as well as percentage of each or both markers that express *Egfr^EM^* (%Marker). (**D**) Estimated densities of SOX9 or OLIG2 (or both) coexpressed with *Egfr^EM^* cells across perinatal ages as indicated (values on y-axis are estimated numbers of cells per mm^3^ × 10^6^). Data in both C and D are mean ± SEM; n=3, individual dots are for each n. Asterisks, Student’s t-test, p < 0.05.

To conduct a thorough examination of markers for progenitor populations and cell cycle regulators, we first quantified the proportion of cycling progenitors that express EGFR during the period of neurogenesis-to-gliogenesis switch and early postnatal forebrain development. All quantifications were analyzed to obtain fractions of all EGFR+ and marker+ cells that overlapped with each other (**Fig. S2**). Bromodeoxyuridine (BrdU) pulse labeling was combined with immunostaining for the mitotic marker phospho-histone H3 (PH3) and EGFR to capture large proportions of actively dividing progenitors at P0 and P30. While a substantial fraction of BrdU^+^ and PH3^+^ progenitors overlap with EGFR during the peak of its expression in the postnatal forebrain, a considerable number of cycling cells were EGFR-negative at any given timepoint. In addition, ratios of EGFR’s coexpression with various established progenitor markers in the SEZ and WM revealed that substantial percentages of EGFR+ cells coexpressed the basic helix-loop-helix Oligodendrocyte transcription factors OLIG1, OLIG2, the Sry-box (SOX9), and Paired box (PAX6) transcription factors at P0 and P30 (**Fig. S2**). EGFR’s overlap was also assessed with the Nuclear Factor IA (NFIA), which is critical for astrocyte development ^20^. We found that NFIA largely overlapped with SOX9 in the SEZ/WM layer, and thus excluded it from subsequent analyses (**Fig. S2**). Some EGFR+ cells coexpressed the oligodendrocyte precursor cell (OPC) marker CC1 [recently identified as a *Qk* gene product ^40^] and radial glia marker SLC1A3 (also known as GLAST), although not as robustly as with the aforementioned markers at both developmental timepoints. Surprisingly, at P0 there was little overlap of EGFR+ cells with other well-established OPC and glial markers expressing the alpha subunit of the platelet derived growth factor receptor (PDGFRα), and Neuron-Glia antigen 2 (NG2), and none was detected at P30.

Since antibody staining can result in false-positive data due to non-specific binding, we confirmed the specificity of the EGFR antibody employed during the various developmental time points using the knock-in *Egfr^Em^* mouse introduced earlier (**Fig. 1A**). We focused on OLIG2 and SOX9 coexpression with *Egfr^Em^* cells since this combination should capture the majority of oligodendrocyte and astrocyte progenitors as well as postmitotic glial populations (**Fig. 1B**). Confirming our findings from antibody staining for EGFR, a large percentage of *Egfr^Em^* + cells in the SEZ/WM coexpressed both SOX9 and OLIG2 (SOX9/OLIG2) at E18.5 (55±1%) and P5 (55±5%), but dramatically and significantly declined by P30 (**Fig. 1C**). In contrast, *Egfr^Em^* + cells expressing either marker but not the other gradually increased in both density (**Fig 1D**) and percentages between E18.5 (SOX9, 14±3%; OLIG2, 2±1%), P5 (SOX9, 21±2%; OLIG2, 17±2%) and later at P30 (SOX9, 55±12%; OLIG2, 24±5%). These dynamic changes were in conjunction with an initial increase in density of *Egfr^Em^* + cells between E18.5 and P5 followed by a sharp and significant decline in their density by P30 (**Fig. 1D**). These findings suggest a temporal and graded EGFR expression in the unique OLIG2+/SOX9+ population in the SEZ/WM that may constitute a distinct set of bi-potential glial progenitors during perinatal development.

### Effects of conditional Egfr deletion on progenitor proliferation, cell death, and differentiation in the developing cortex

To elucidate the requirement for EGFR during forebrain development, we carried a conditional *Egfr* allele (*Egfr^F/F^*) which was used in two of our previous studies ^18,34^, onto a transgenic *Nestin-cre* background to target both the ventral and dorsal telencephalic progenitors (*Nes:F/F;***Fig. 2A**). Additionally, *Egfr^F/F^* mice were bred with *Emx1^cre^* mice to restrict deletion to dorsal telencephalic progenitors (*Emx:F/F;* **Fig. 2A**)^41^. Robust deletion of EGFR in the forebrain was confirmed in *Nes:F/F* brains by PCR, Western blotting, and immunohistochemistry (**Fig. S3**). Despite the normal appearance of *Nes:F/F* pups at birth (P0), defects became apparent at P7 with spots of hemorrhage scattered within the rostral forebrain (**Fig. 2B**) along with significantly smaller body size and decreased weight gain between P0 and P56 (**Fig. S3**). In contrast, *Emx:F/F* mice did not exhibit developmental defects as their forebrains appeared grossly similar to *WT* controls (*WT*; **Figs. 2B–2C**), suggesting that dorsal progenitors appear to be EGFR-independent at the bulk-level of analysis. Alternatively, EGFR-dependent progenitors from outside the dorsal telencephalon (e.g., EGFR+ progenitors in the ventral forebrain) may compensate for the missing EGFR-dependent glia in the *Emx:F/F* forebrain. For this reason, we largely focused on characterizing the clear defects in the *Nes:F/F* forebrain for the remainder of our bulk analyses. In addition, no obvious gross morphological or noticeable cellular differences were found in *Nes:F/+* or *Emx:F/+* forebrains, indicating that heterozygous deletions of *Egfr* fail to result in obvious defects.

**Figure 2.**
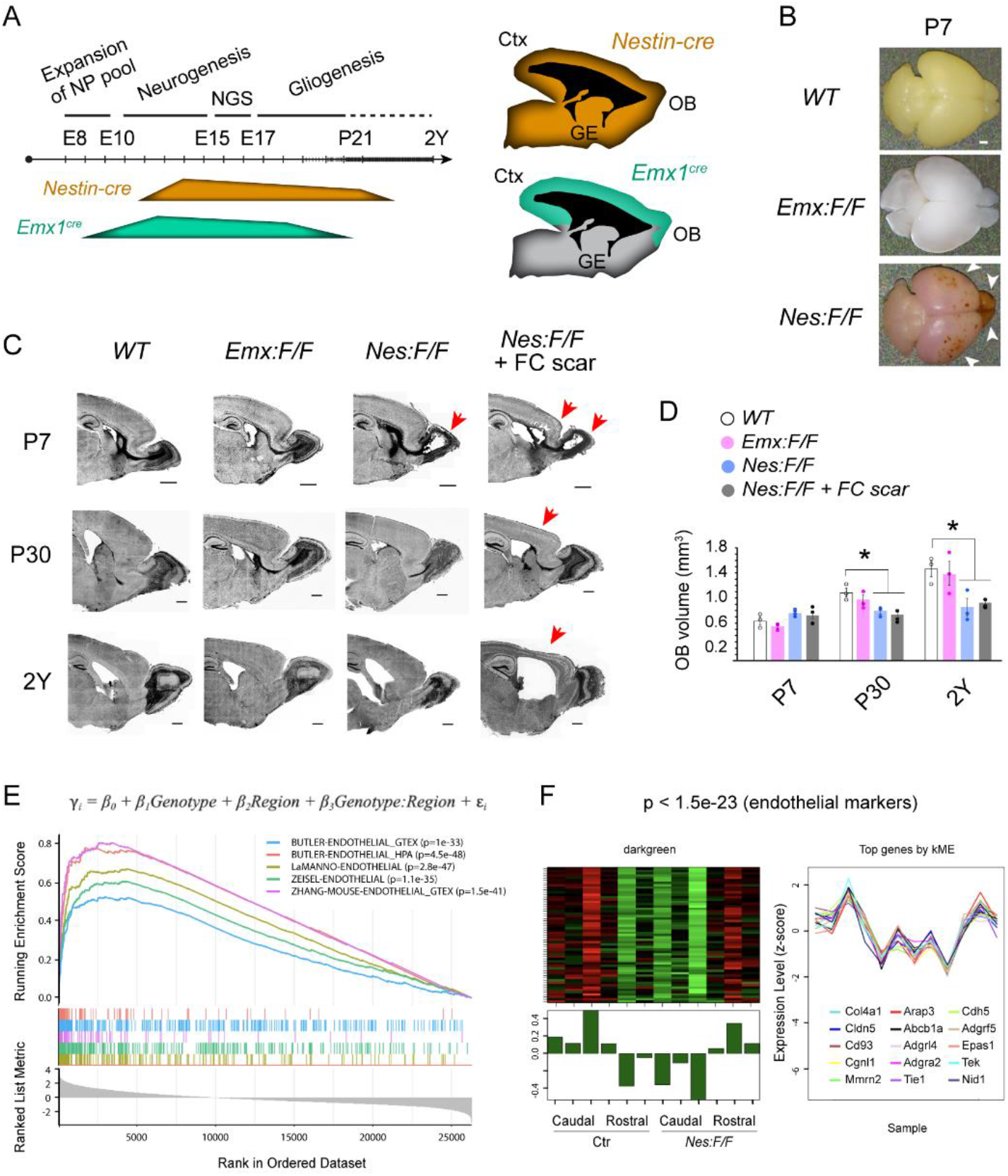
Gross defects in the *Nes:F/F* forebrains during early postnatal development and aging. **(A)** Differences in temporal and spatial expression of cre in *Nestin-cre* and *Emx1^cre^* mice allows for capturing effects of *Egfr* deletion in distinct progenitor populations of the forebrain as describe before ^41^. **(B)** Dorsal view of dissected *WT*, *Nes:F/F* and *Emx:F/F* brains at P7. Hemorrhage and blood accumulation is evident in the frontal cortices and olfactory bulbs of the *Nes:F/F* mice (arrowheads). Pink tint in the *Nes:F/F* brain is due to presence of tdTomato cre recombinase reporter. Scale bar, 1 mm. **(C)** DAPI stained sections of forebrains at P7, P30, and 2Y for each genotype and in *Nes:F/F* mice exhibiting cortical injury. Arrows point to regions with scarring or altered histology such as the cavity in the OB at P7 which disappears by P30. Scale bars, 500 μm. **(D)** Olfactory bulb volumes of *WT*, *Emx:F/F*, *Nes:F/F*, and *Nes:F/F* mice with frontal cortical scar (FC scar) and mice across various ages. Data are mean ± SEM; individual dots are for each n. Asterisks, Student’s t-test, p < 0.05. **(E)** Transcriptional changes in E17.5 *Nes:F/F* forebrain samples were analyzed via multiple linear regression. Chart illustrates genes in descending order for the regression coefficient of the interaction term used as input for gene set enrichment analysis (GSEA). Genes upregulated in rostral *Nes:F/F* samples were significantly enriched for multiple endothelial gene sets. **(F)**A gene coexpression module enriched for endothelial markers is upregulated in rostral vs. caudal samples from control (Ctr) and *Nes:F/F* mice at E17.5. The heatmap indicates the relative expression levels of the module genes across all samples, and the module eigengene (first principal component) below summarizes its expression pattern. Right plot, z-scored expression levels of the top 15 genes correlated with the module eigengene.

The hemorrhages at P7 were observed on the surfaces of all *Nes:F/F* OBs, while only 45% of these mice (n=20) showed additional spots in the frontal cortices. Histological analyses revealed that tissue disruption was always present in the *Nes:F/F* OB, including damage to the granule and mitral cell layers and presence of a large cavity at the core of the OB as early as P7 (**Fig. 2C, arrow**). The cavity corresponds to the earlier position of the olfactory ventricle during embryonic and early postnatal development, which closes prior to P7 in mice ^42,43^. Curiously, the gross defects in the *Nes:F/F* OB appeared resolved at P30 (**Fig. 2C**), yet the size of the OB was significantly smaller than *WT* and *Emx:F/F* OBs at P30. The *Nes:F/F* OB remained smaller in mice with and without frontal cortical scarring up to 2 years (2Y; **Fig. 2D**). Moreover, cortical scarring corresponded with signs of hemorrhage at earlier stages, suggesting that the damage was resolved in a similar manner to that observed in the OB. However, 19% of the *Nes:F/F* mice (n=21) showed persistent scarring in the frontal cortices at P30, suggesting that unlike the OB, the frontal cortex was more susceptible to long-term damage in response to the hemorrhaging during early postnatal development. To better understand the onset of necrosis, bleeding and scarring in the rostral *Nes:F/F* cortices, we dissected rostral and caudal halves containing cortical areas and the hippocampal formation from E17.5 embryos. Samples were subjected to RNA-sequencing and bioinformatics analyses revealing robust upregulation of endothelial gene sets ^15,44–46^ in rostral, but not caudal cortices of *Nes:F/F* forebrains (**Figs. 2E–2F**). This finding likely reflects a vascularization response to pre-scarring changes in the frontal half of the forebrain that may be related to defective gliogenesis in the affected tissue. In summary, the perinatal *Nes:F/F* bleeding and scarring in the frontal forebrain appear to be dispensable for survival of mice (unlike germline deletion of *Egfr*), coincident with resolution of most histological anomalies that emerge during the first week of postnatal life in the *Nes:F/F* forebrain. Based on the timing of EGFR expression, emergence of perinatal defects in vascular homeostasis, and resolution of the anomalies by P30, we reasoned that late neurogenesis and/or gliogenesis may be compromised in the rostral *Nes:F/F* forebrain.

To assess the extent of EGFR’s impact on progenitor homeostasis, we quantitatively compared the proliferation rate of *WT* and *Nes:F/F* progenitors using acute BrdU pulsing *in vivo* which labels cells undergoing DNA replication or repair (**Fig. S4**). While the density of BrdU labeled cells at P7 was significantly lower in *Nes:F/F* than in the *WT* SEZ/WM (encompassing both ventral and dorsal regions), they were surprisingly similar to *WT* levels at P0 or P30. This suggested that either EGFR plays a temporally transient role in progenitor proliferation, or that it is required for expansion of a distinct population of progenitors during the first week of postnatal life. Further assessment of expansion capacity of progenitors extracted from the forebrain using the neurosphere assay ^47^ revealed a transient defect whereby the size, but not number of neurospheres, was significantly smaller in primary *Nes:F/F* cultures than in *WT* controls suggesting impaired replication capacity or cell death (**Fig. S4**). Interestingly, both parameters were similar to *WT*-derived neurospheres after the fourth passage, matching the recovery observed *in vivo*.

To evaluate if the lower number of proliferating cells was due to defects in cell division or cell survival, we quantified levels of cleaved caspase3 (c-CASP3) immunoreactivity as a marker for apoptosis in the forebrain (**Fig. S5**). Estimates of c-CASP3^+^ cell densities were indistinguishable between the *WT* and *Nes:F/F* SEZ/WM from P0 to P30, similar to the OB at P0 (**Fig. S5**). However, we readily found c-CASP3^+^ particles and cells around the damaged area in the *Nes:F/F* OB and cortices, indicating significant levels of apoptosis. The damage in the *Nes:F/F* OB appeared repaired by P30, accompanied with reduction in densities of c-CASP3^+^ cells to *WT* levels. As mentioned earlier, the scar in the fraction of cortices with perinatal damage failed to resolve throughout life. Combined, our *in vivo* and *in vitro* analyses of *Egfr* deletion in the forebrain indicate that EGFR is critical for a developmentally transient and/or subpopulation-specific expansion of ventrally derived forebrain progenitors during the first postnatal week in mice. However, overall proliferation of progenitors was largely compensated by the third postnatal week in *Nes:F/F* forebrains, presumably by progenitors that are EGFR-independent. Moreover, the gross phenotypes of *Egfr* deletion are only visible when both dorsal and ventral domains of the forebrain are targeted, whereas dorsal deletion alone fails to cause any obvious defects in terms of general forebrain development and homeostasis.

Based on the known associations between EGFR expression and glial production, we next quantified the same series of markers analyzed in the first part of the study, in the *Nes:F/F* SEZ/WM during early postnatal periods (**Fig. S6**). The most significant effect was seen in depleted densities of populations of OPCs expressing OLIG2 at all ages, and OLIG1 at P7 and P30 using immunohistochemistry (**Fig. S6**), which was confirmed in RNA-seq results obtained from E17.5 *Nes:F/F* cortices (**Fig. 3A**). RNA-seq analyses of rostral and caudal cortices at P5 showed that depletions of EGFR, OLIG1 and OLIG2 were maintained in *Nes:F/F* forebrains (**Fig. 3A**). In addition, altered expression of astrocytes became evident at this early postnatal stage indicating disruptions in both astrocyte and oligodendrocyte populations in the absence of *Egfr* in the rostral forebrain by P5 (**Fig 3B**). Substantial upregulation of myeloid, microglial and macrophage transcripts correlated with the histological necrosis noted in the rostral cortices at this age (**Fig. S7**).

**Figure 3.**
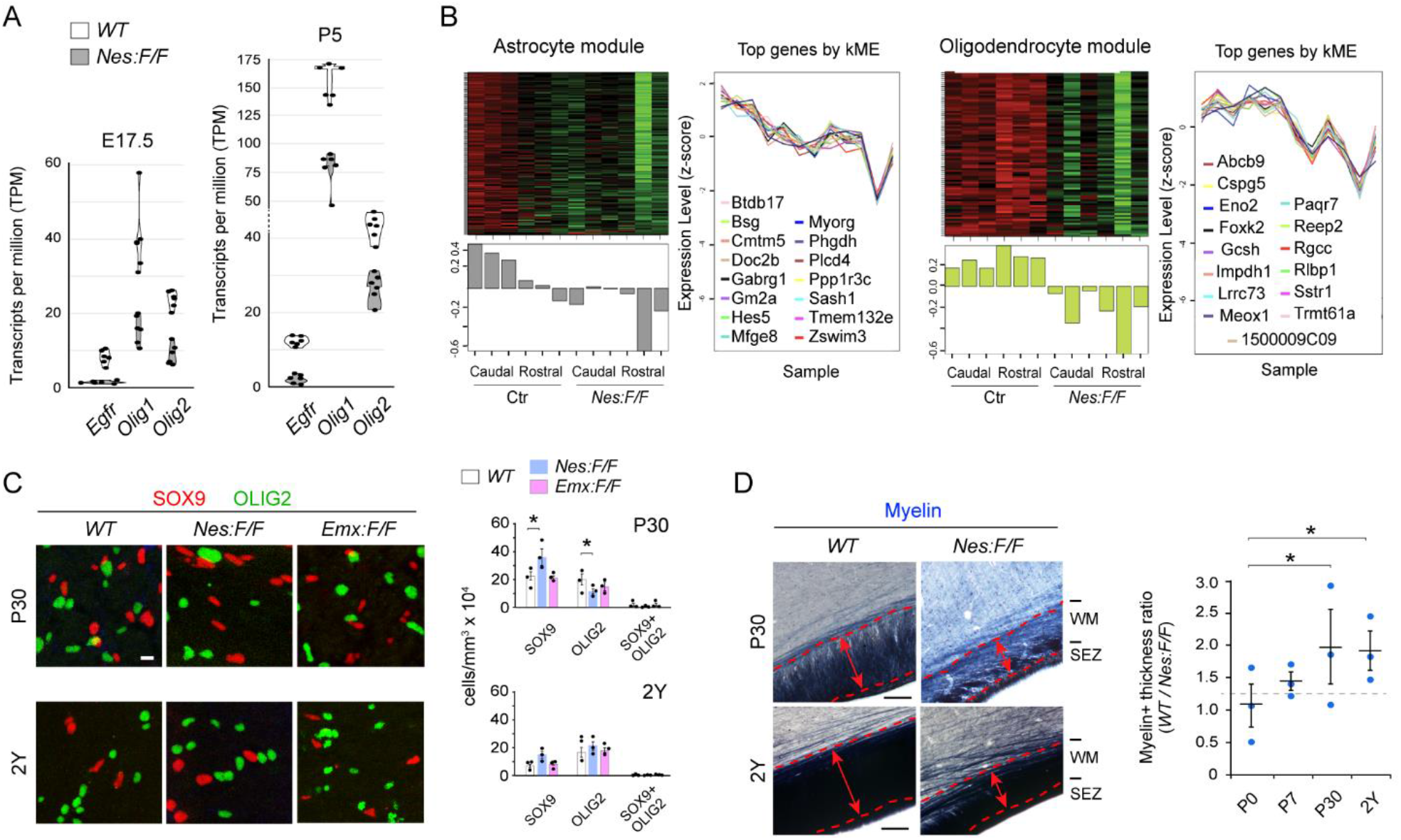
Characterization of gliogenesis and myelin content in the cortex over the life-span of *Nes:F/F* mice. **(A)** RNA-seq comparison of cortical samples obtained from E17.5 embryos and P5 neonates indicate that *Egfr*, *Olig1*, and *Olig2* transcripts were significantly reduced in *Nes:F/F* cortices compared to *WT* controls. TPM, transcripts per million. **(B)** Gene coexpression modules enriched for Astrocyte and Oligodendrocyte markers reveal differential expression in rostral vs. caudal samples from control (Ctr) and *Nes:F/F* mice at P5. Heatmaps illustrate the relative expression levels of module genes across all samples. The module eigengene (first principal component) below summarizes its expression pattern. Right plots, z-scored expression levels of the top 15 genes correlated with each module eigengene. **(C)** Confocal images of cortical SEZ/WM regions of *WT*, *Nes:F/F*, and *Emx:F/F* mice at P30 and 2Y. Scale bar, 10 μm. Bar chart represents mean densities of each marker ± SEM; dots are values for each n. Asterisks, Student’s t-test, p < 0.05. **(D)** Myelin stained SEZ/WM of *WT* and *Nes:F/F* mice at P30. Dotted lines delineate the WM, the thickness of which is indicated by the double arrows. Bar chart, mean myelin thickness ± SEM; dots are values for each n. Asterisks, Student’s t-test, p < 0.05.

In vivo analysis of estimated densities for established markers in the SEZ/WM using antibody staining revealed a temporal gradient of alterations in control and *Nes:F/F* cortices (**Fig. S6**). For example, while cells positive for the OPC marker PDGFRα were significantly reduced early postnatally, SOX9 and S100 (radial glial and astrocyte markers) were significantly less abundant at P7 but not the other ages. In fact, density of SOX9+ cells was significantly elevated in the *Nes:F/F* SEZ/WM, suggesting compensation and potential over production of astrocytes and/or radial glial-like cells at P30. Interestingly, PAX6, which is largely known as a neurogenic progenitor marker ^48^, was selectively and significantly upregulated at P7, suggesting potential elevated neurogenesis in the *Nes:F/F* SEZ/WM, which recovered to *WT*-levels by P30 (**Fig. S6**). Significant decline in densities of other OPC markers OLIG1, OLIG2, and NG2, were evident at P7 and P30. Other markers were reduced significantly at P30 but not earlier, which included SLC1A3, CC1, and the pan-DNA label DAPI (**Fig. S6**). Collectively, the characterization of known markers in the perinatal SEZ/WM suggests that EGFR is required for generation and maintenance of significant fractions of OLIG1/2+ OPCs during early postnatal development. Despite the absence of EGFR, gliogenesis appears to be able to partially recover by P30. If this is through an EGFR-independent mechanism or compensation by a foreign population of EGFR+ progenitors remained unclear at this juncture.

Since a significant portion of EGFR+ cells coexpressed SOX9 and OLIG2 in our earlier analyses, we next opted to simultaneously visualize the two transcription factors in control and *Nes:F/F* mice through early developmental stages and later during aging (**Fig. 3C; Fig. S8**). Both markers report on astrocyte and oligodendrocyte progenitors and their postmitotic populations ^10,49–51^, but little is known regarding their overlap during the developmental gliogenic periods. Focusing on neocortical areas, the distribution of the two markers in the SEZ/WM, the deep (DL), and upper (UL) layers of the *WT* cortex, revealed spatial and temporal gradients for cells that coexpress SOX9 and OLIG2, or each marker exclusively. Interestingly, most populations were distinctly labeled by one marker but not the other in the SEZ/WM of early postnatal mice between P0 to P30. However, there were substantial numbers of SOX9 cells colabeled with OLIG2 in the DL and UL of dorsolateral cortices at P0 and P7, but not at P30 (**Fig. S8**).

In the *Nes:F/F* cortices, there was a significant depletion of SOX9, OLIG2, and OLIG2/SOX9 positive cells at P0 in all layers of the cortex, but the densities remarkably recovered for all three populations in the UL and DL by P7 **(Fig. S8)**. Interestingly, the density of SOX9+ population, which constitutes astrocytes, was significantly elevated in all three layers of the cortex at P30. This elevation may at least partially include reactive and scar forming population of astrocytes that had responded to the perinatal damage described earlier in the rostral forebrain. In contrast, the density of the OLIG2+ population was lower in the SEZ/WM, while at the same level as control mice in the UL and DL layers (**Fig. S8**). Though the elevated SOX9 population was still evident in the 2Y SEZ/WM, the OLIG2+ population was no longer significantly different than controls in the aged cortex (**Fig. 3C**). Additional assessment of for the same markers indicated an initial change in distribution of SOX9, OLIG2, and SOX9/OLIG2 in UL and DL of *Emx:F/F* cortices at P0, but not in the SEZ/WM (**Fig. S8**). These changes normalized by P7 (**Fig. S8**) and were indistinguishable from controls by P30 and later at 2Y (**Fig. 3C**). The elevated SOX9+ cells in *Nes:F/F* cortices were not found in *Emx:F/F* mice (**Fig. 3C**) confirming that the gliogenic phenotypes identified require inclusion of the Nestin domain that spans both dorsal and ventral aspects of the forebrain. Together our findings further substantiate the possibility that the OLIG2/SOX9 double positive cells identified in our earlier characterization of EGFR expression (**Fig. 1B**) may function as bipotential glial progenitors under the regulation of EGFR signaling.

Since overall cell proliferation, apoptosis, and production of astrocytes appeared normalized by P30 in the *Nes:F/F* forebrain, selective depletion of key OPC markers at P0 and P30 suggested that the observed gross defects may be due to faulty myelination in the forebrain. To assess this, we stained for myelin at P0, P7, P30, and 2Y in *WT* and *Nes:F/F* forebrains (**Fig. 3D**). The myelin+ WM was significantly thinner in *Nes:F/F* mice than *WT* controls beginning at P7, which persisted until 2Y (**Fig. 3D**). However, some myelination appeared preserved in the *Nes:F/F* forebrains based on the consistency of staining in the WM after P30. The residual myelin could be due to compensation by alternative mechanisms, or possible reemergence of EGFR expression from ‘escaper’ cells due to reported incomplete deletion inherent to the Cre-Lox system ^52^. To determine if any escaper cells were capable of repopulating the EGFR-depleted pool of progenitors during the life span of *Nes:F/F* mice, we labeled SEZ sections at P7, P30, or 2Y with the EGFR antibody and found little to no EGFR immunoreactivity in the *Nes:F/F* SEZ and WM compared to controls (**Fig. S9**). Thus, the apparent developmental compensation for loss of EGFR seems independent of any *de novo* EGFR expression from potential cells that escape cre-mediated recombination. Taken together, our findings indicate that myelination occurred to some extent in the absence of EGFR-dependent oligodendrocyte production in the cortex. While it is possible that EGFR-dependent mechanisms are normally responsible for all myelination, some mechanisms that normally use EGFR signaling can still occur in the absence of EGFR (possibly in a new, compensatory way).

### Reactive gliosis reported by GFAP expression is preserved in the absence of Egfr

To directly assess the subpopulations of reactive and SEZ/WM astrocytes, we used antibody staining against glial fibrillary acidic protein (GFAP) to measure its expression in *WT* and *Nes:F/F* forebrains. We split the analysis into rostral and caudal halves of the cortices in sagittal sections since we had earlier found rostral cortical scarring in some *Nes:F/F* mice. Regardless of the presence of scarring in the frontal cortices, there was no significant change in GFAP intensity in the SEZ/WM, or neuronal layers of the caudal cortices in *Nes:F/F* mice compared to controls at P0 or P30 (**Fig 4A**). However, there was a significant increase in GFAP intensity in all layers of caudal cortices in P7 *Nes:F/F* mice with frontal cortex scarring (+ FC scar; **Figs. 4A, 4B**). In frontal cortices of mice with FC scar there was a significant elevation of GFAP labeling at both P7 and P30 in all cortical layers (**Fig. 4A, 4B**). Since reactive glia respond to damage in the CNS and are critical for formation of a scar, the increase in GFAP+ cells likely constitutes a secondary effect to the transient tissue damage in the *Nes:F/F* forebrain. To test this possibility further, we inflicted blunt cortical stab injuries in P0, P30, and 2Y *WT* and *Nes:F/F* frontal cortices followed by a one-week recovery period (**Fig. 4C, 4D**). While levels of GFAP immunoreactivity were significantly lower in P0 stab-injured cortices, GFAP+ gliosis was observed around the site of injury with no quantitative differences between *WT* or *Nes:F/F* cortices (**Fig. 4D**). These findings supported the hypothesis that reactive gliosis and WM astrocytes are at least in part independent of EGFR expression.

**Figure 4.**
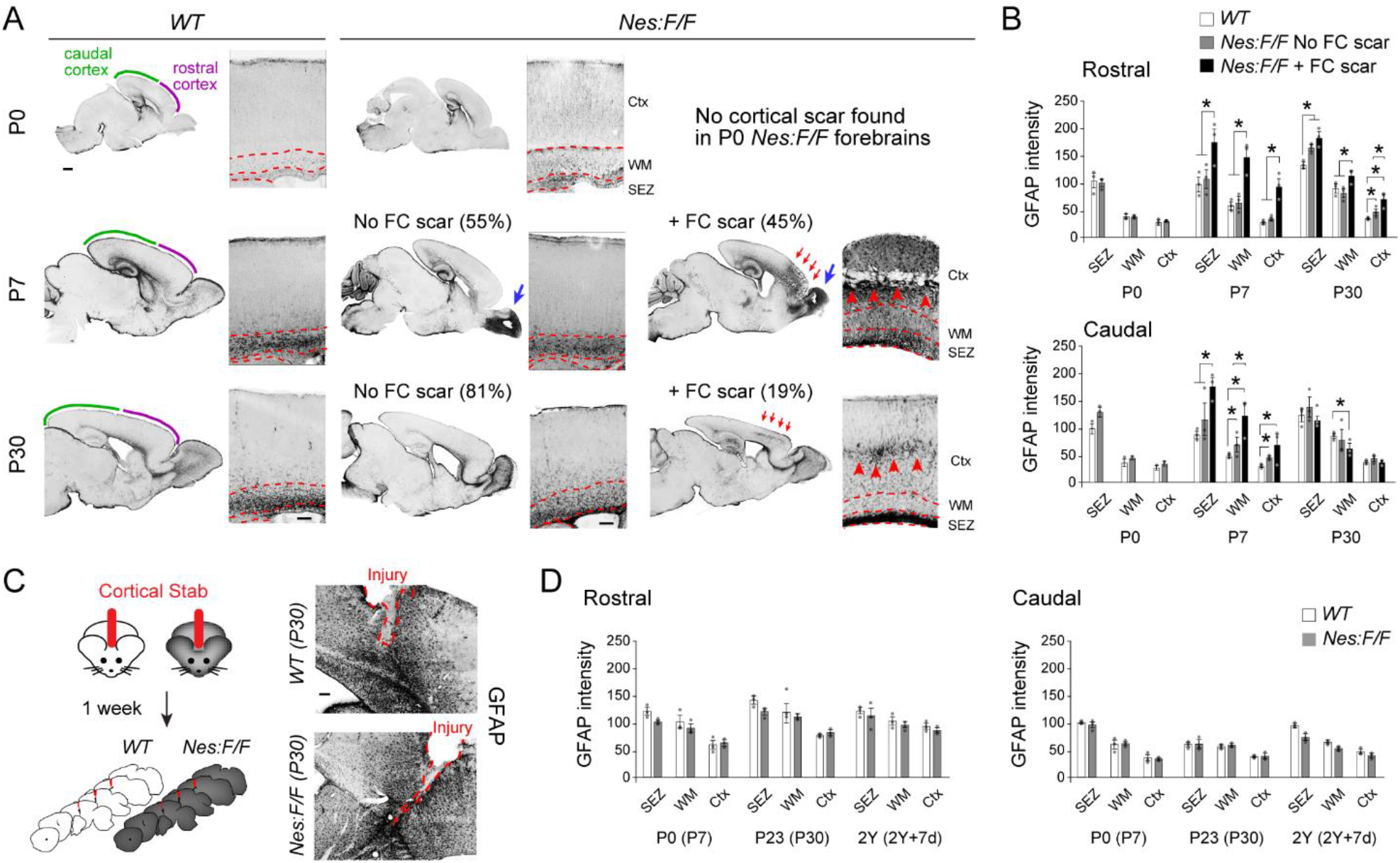
Characterization of WM and reactive astrocytes in the *Nes:F/F* forebrain. **(A)** Confocal images of GFAP stained frontal cortices (Ctx) at P0, P7, and P30 in *WT* and *Nes:F/F* cortices. Dotted lines demarcate boundaries between SEZ, WM, and Ctx; Necrosis/injury is observed in *Nes:F/F* OBs at P7 (large blue arrow). Some *Nes:F/F* mice exhibit no frontal cortical injury (No FC scar) while others have scarring in the frontal cortices (+ FC scar) at P7 and P30 (small red arrows). Percentages in parentheses are for each phenotype as indicated (n=20, P0; n=20, P7; n=21, P30 *Nes:F/F* mice). **(B)** Quantification of GFAP fluorescence intensity in each region across the three ages in *WT* and *Nes:F/F* forebrains with and without FC scarring. **(C)** Schemata of cortical stab injuries followed by tissue analysis a week later. Confocal images of the *WT* and *Nes:F/F* injured cortices (red dotted line) stained for GFAP (dark signal). **(D)** GFAP intensity quantified in cortices containing the stab injury (age of stab injury, age of analysis in parentheses). Data in B and D are mean ± SEM of immunofluorescence intensity, n=3 animals/genotype and age. Asterisks, Student’s t-test, p < 0.05. Scale bars, 100 μm.

Finally, to assess neuronal integrity in *Nes:F/F* cortices we conducted NeuN immunostaining at P0 and P30 to visualize mature neurons in layers of the cortex (**Fig. S10**). Results showed no alterations in density of NeuN+ cells in rostral cortical areas where necrosis was at least partially present. Moreover, analysis of layer-specific cortical markers CTIP2 and CUX1 indicated no measurable differences in the layering of these markers in the *Nes:F/F* frontal cortices (**Fig. S10**). Taken together these findings further support the hypothesis that the loss of EGFR impacts gliogenesis but not neurogenesis during forebrain development in mice.

### Sparse MADM-based deletions of Egfr reveal the cell autonomous role in gliogenesis but not neurogenesis in the forebrain

Concerned that our analysis of neuronal integrity in *Nes:F/F* mice may be hampered by non-autonomous effects of gliogenesis on cells in the cortex, we opted to utilize alleles for Mosaic Analysis with Double Markers (MADM) ^53^ which allow for sparse deletion of *Egfr* in forebrain progenitors in combination with permanent fluorescent labeling of cells undergoing recombination. To achieve this, we employed MADM alleles on the mouse chromosome 11 (MADM-11) ^54^, where the *Egfr* locus is also localized. MADM operates through two reciprocal chimeric reporter genes inserted near centromeric regions of chromosomes. Cre-mediated recombination of these chimeric reporter genes during mitosis, with one allele co-carrying a single allele of the *Egfr floxed* domain, results in definitive labeling of cells carrying *WT* (tdTomato^+^, red), heterozygous (tdTomato^+^/GFP^+^, yellow), and homozygous (GFP^+^, green) *floxed* alleles of *Egfr* at low densities (*MADM:F/*+ mice; **Fig. S11**). Since the MADM recombination events result in permanent labeling of the daughter cell and its corresponding lineage, we can unambiguously track genetically manipulated cells. Furthermore, because genetic reporters in *MADM:F/*+ tissues arise from cre-mediated recombination in rare (~1:1000) mitotically recombining progenitors, fate specification is tracked in clonally derived *WT* and *Egfr-null* sibling daughter cells ^18,34^. MADM-11 animals that do not carry a *floxed* allele for *Egfr* were used as controls (*MADM:*+/+ mice; all three combinations of reporter^+^ cells are *WT).* In addition, mice carrying both *floxed Egfr* alleles were used as in the bulk analysis described earlier, allowing for analysis of MADM labeled lineages in the bulk *null* background which contained scarring in the rostral forebrain (*MADM:F/F* mice; all three combinations of reporter^+^ cells are *Egfr-null).* The same two cre-drivers used in the earlier describe bulk analyses were employed; *Emx1^cre^* to restrict cre-mediated recombination to the dorsal telencephalon (*Emx:MADM:*+/+, *Emx:MADM:F/*+, and *Emx:MADM:F/F*) and *Nestin-cre* to drive expression throughout the CNS (*Nes:MADM:*+/+, *Nes:MADM:F/*+, and *Nes:MADM:F/F;* **Fig. S11**). To determine if mosaic presence of *Egfr-null cells* in *Nes:MADM:F/*+ and *Emx1:MADM:F/*+ forebrains results in the necrosis phenotype observed at P7 in the *Nes:F/F* bulk deletion model, we performed GFAP staining. Unlike bulk-deleted *Nes:F/F* forebrains, *Nes:MADM:F/*+, *Emx:MADM:F/*+, and *Emx:MADM:F/F* mice failed to exhibit tissue damage or reactive gliosis in the rostral forebrain (**Fig. S11**). In line with the earlier bulk results, *Nes:MADM:F/F* forebrains exhibited the same tissue damage as *Nes:F/F* mice.

It became immediately and qualitatively obvious that *Nestin-* and *Emx1-*driven MADM recombinations resulted in distinct and surprising phenotypes (**Fig. 5A**). Since we observed a rostral forebrain defect upon bulk *Egfr* deletion in *Nes:F/F* mice, we divided our quantitative MADM analyses into rostral and caudal domains of the cerebral cortices. As expected, MADM cells were present in both dorsal and ventral regions of the *Nes:MADM:*+/+ forebrain, whereas *Emx1:MADM:*+/+ cells were dorsally restricted to the cortex and the hippocampal formation (**Fig 5A, Movies S1-S6**). In both control MADM forebrains, neurons and glia were observed in all cortical layers (**Fig. 5**), as well as various cell types within the SEZ/WM (**Fig. S12**). A striking phenotype emerged upon introduction of a single floxed *Egfr* allele to generate clones in which red (tdTomato) cells are *Egfr* ^+/+^ and green (GFP) cells are *Egfr* ^-/-^ on a largely *Egfr* ^+/-^ cellular backgrounds. In both *Emx:MADM:F/*+ and *Nes:MADM:F/*+ cortices, green and red neurons were present, but nearly all MADM-labeled glia and cells in the SEZ/WM were red (i.e., *WT).* Detailed characterization of various cell types labeled by both *Nestin* and *Emx1* cre drivers used in our study (**Fig. 5B**) indicated that the expanded red glia largely comprised astrocytes in the UL and DL (neuronal layers) of the *F/*+ cortices (**Fig. 5C**).

**Figure 5.**
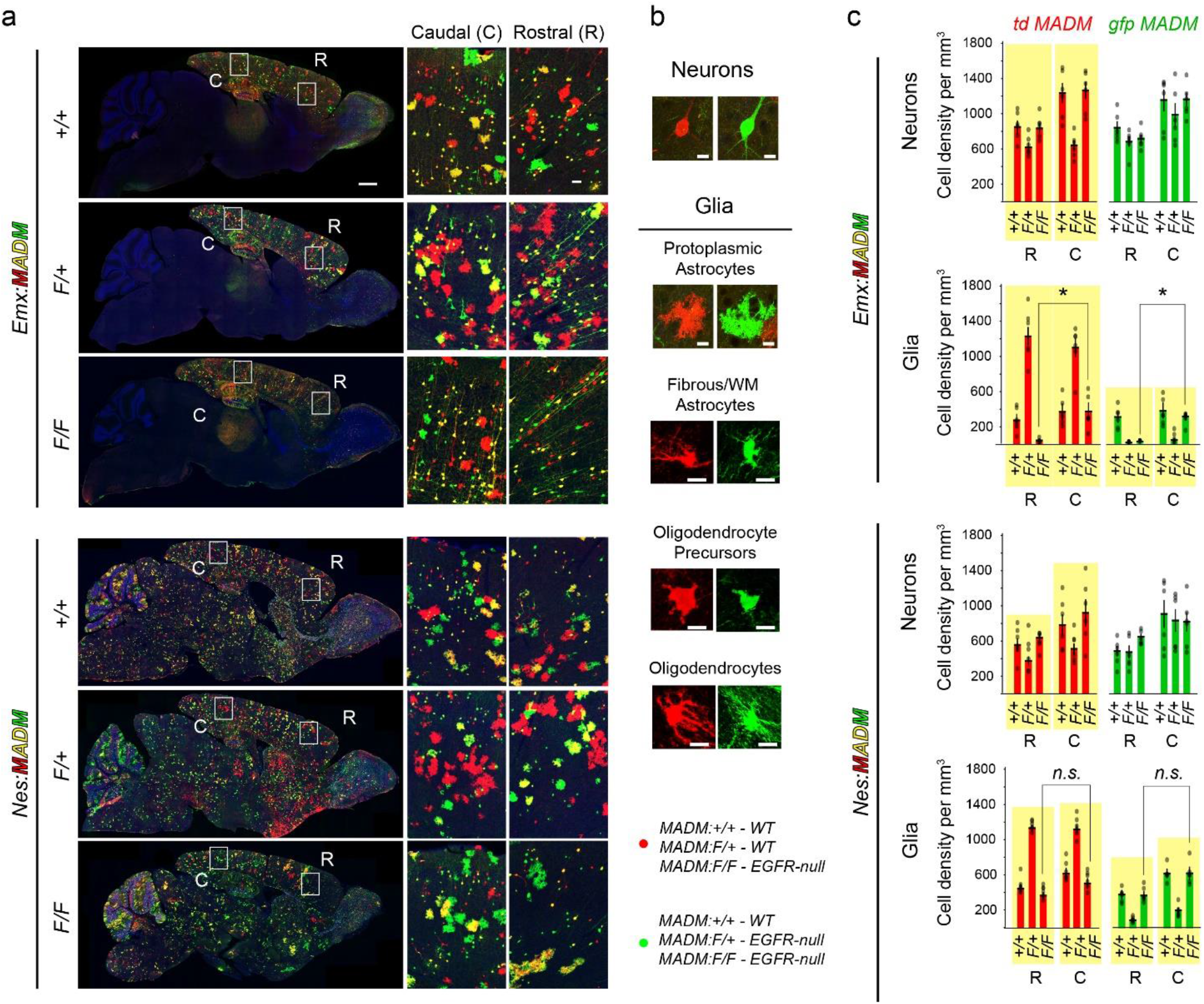
Regional and cellular phenotypes associated with *Egfr* genotypes in upper and deep (neuronal) layers of *MADM* cortices. **(A)** Representative confocal images of P30 *Emx:MADM* and *Nes:MADM* forebrains with different *Egfr* genotypes as indicated. Sample high magnification images used for morphological characterization of distinct green and red neurons and glial types (right two columns in each image). Scale bars, sagittal views, 1 mm; zoomed panels in caudal and rostral areas, 50 μm. **(B)** Morphology-based characterization of neurons and glia (oligodendrocytes and astrocytes) at high magnification. Some pericytes were also labeled in *Nes:MADM* lines, but their numbers were extremely low compared to other cell types. Scale bars, 10 μm. **(C)** Rostral and caudal cell densities of MADM labeled neurons, glia, and pericytes in the neuronal cortical layers (see Figs. 6 and S12 for SEZ/WM characterizations and data). N=6 mice per genotype. Asterisks, Pairwise or Student’s t-test, p < 0.05. White arrows in A and black arrow in C, point to elevated MADM astrocytes in caudal *Emx:MADM:F/F* cortices relative to rostral where glia are nearly abolished.

Surprisingly, *Nestin* and *Emx1* cre-driven recombination in *MADM:F/F* cortices resulted in distinct phenotypes. While red and green MADM glial densities in *Nes:MADM:F/F* mice appeared similar to those in control rostral and caudal cortices, red and green MADM lineages in *Emx:MADM:F/F* cortices contained almost no glia in rostral cortical areas (**Fig. 5A, 5C**). In contrast, some red and green *MADM* glia reemerged in caudal *Emx:MADM:F/F* cortical layers (**Fig. 5A, 5C, arrows**). This data is suggestive of a gradient of EGFR-dependent and EGFR-independent progenitors along the rostrocaudal and dorsoventral extents of the forebrain. Accordingly, a rostro-dorsal gliogenic progenitor population appears to be highly EGFR-dependent, whereas an EGFR-independent gliogenic population appears to be situated in a ventrocaudal domain of the forebrain. This population overlaps with both the *Emx1* and *Nestin* progenitor populations.

Since most of the progenitor proliferation occurs in the SEZ/WM layer of the cortex, we next conducted detailed analyses of its rostral to caudal extent in all MADM genotypes (**Fig. 6A**). As expected the MADM-labeled cell types in both *Emx:MADM* and *Nes:MADM* SEZ/WM were more complex than in the UL and DL despite lacking neurons (**Fig. 6B**). However, like the neuronal layers, red MADM OPCs and fibrous astrocytes were highly expanded in the *F/*+ SEZ/WM of both groups of mice **(Figs. 6A, 6C, S12)**. The other cell types in the *F/*+ SEZ/WM were low in density and the effect of *Egfr* deletion was less obvious on these populations. In the *F/F* genotype which constitute the bulk models, oligodendrocyte, fibrous astrocyte, and radial glia populations were all significantly depleted in both *Emx:MADM* and *Nes:MADM* SEZs (**Figs. 6C; S12**).

**Figure 6.**
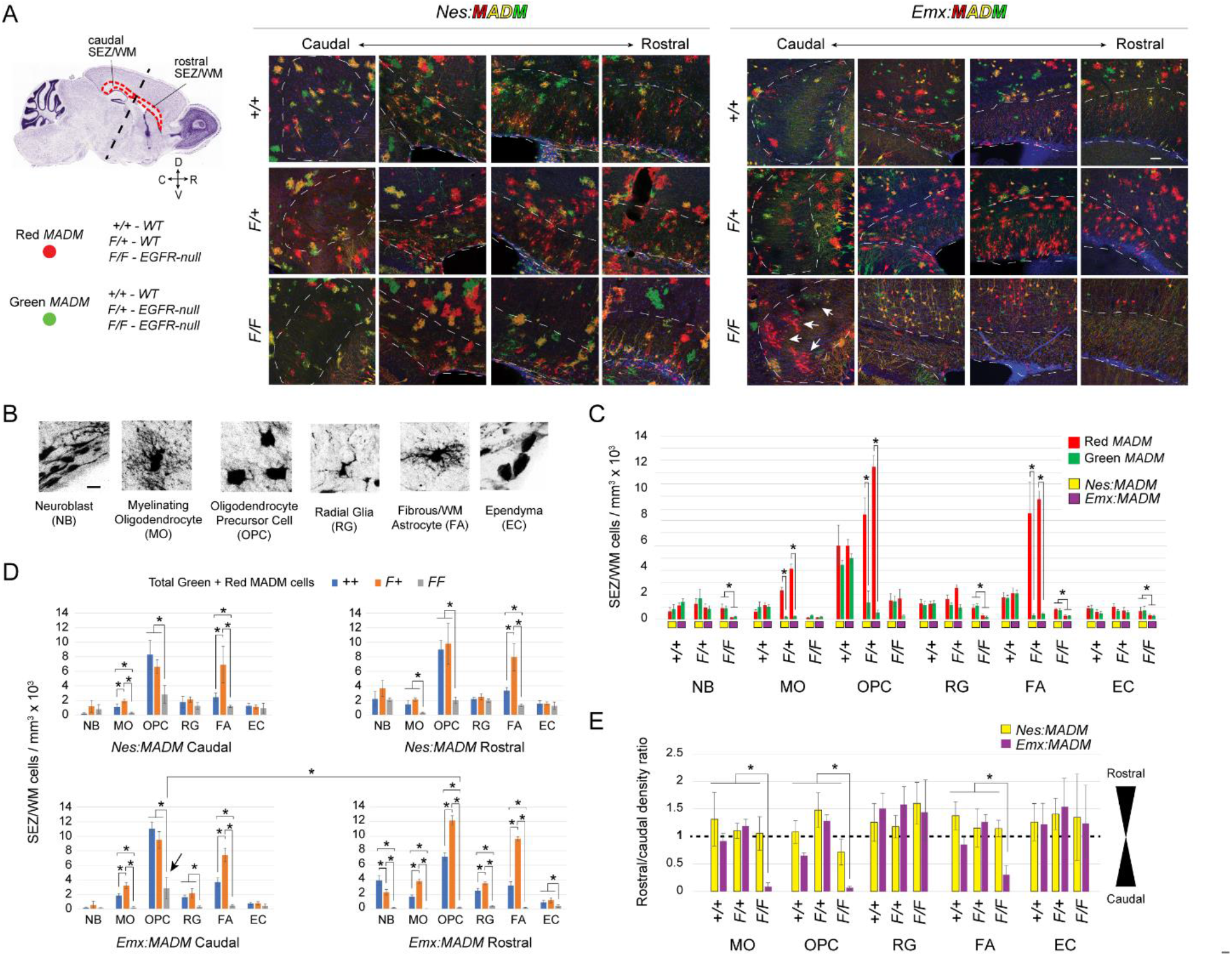
Analysis of cell types in the SEZ/WM of *MADM* forebrains. (**A**) Left, Nissl-stained sagittal section of the mouse brain with dotted areas demarcating rostral and caudal SEZ/WM tissue analyzed for MADM cell types. Red and Green MADM cells possess distinct genotypes as indicated. Right, Confocal micrographs of SEZ/WM (dotted lines demarcate the territories) along rostral and caudal extents of various MADM genotypes as indicated. Scale bar, 50 μm. (**B**) Samples of the MADM cell types found in the SEZ/WM that were used for subsequent characterization. Scale bar, 10 μm. (**C**) Densities of green and red cell types in the SEZ/WM of *Nes:MADM* (yellow boxes) and *Emx:MADM* (purple boxes) forebrains. (**D**) Densities of total MADM cells (Red and green cells combined) in the SEZ/WM for distinct genotypes as indicated. Arrow points to elevated MADM astrocytes in caudal *Emx:MADM:F/F* cortices relative to rostral. (**E**) Ratios of rostral divided by caudal total MADM cell densities (D) for each genotype as indicated. Note, severe depletion of cell densities of oligodendrocytes (MO and OPC) and fibrous astrocytes (FA) in the *Emx:MADM:F/F*, but not *Nes:MADM:F/F* SEZ/WM. Bar charts in C, D, and E are data as mean ± SEM; Asterisks, Student’s t-test, p < 0.05.

A question regarding the striking *F/*+ phenotype is whether the increase in populations of wildtype (red) glia constitutes a compensatory response. We assumed that compensation may be inferred if the combined populations of red and green MADM cells are similar between *F/*+ and +/+ genotypes. In fact, this combination in total SEZ/WM densities revealed that most OPCs and other cell types were similar between *F/*+ and +/+ genotypes (**Fig. 6D**). The exceptions were myelinating oligodendrocytes and fibrous astrocytes in both *Emx:MADM:F/*+ and *Nes:MADM:F/*+ forebrains which had densities beyond the control levels. In addition, OPCs only in the rostral *Emx:MADM:F/*+ domain appeared to have expanded beyond their +/+ densities (**Fig. 6D**). The same analysis on the *F/F* genotypes revealed robust and highly significant depletion of all cell types in the rostral *Emx:MADM* SEZ/WM compared to all other samples (**Fig. 6D**) Curiously, while the oligodendrocyte and astrocyte lineages in the *Nes:MADM:F/F* SEZ/WM remained significantly lower than controls, their densities were raised significantly above the *Emx:MADM* levels (**Figs. 6A, 6D**, arrows). To better visualize this regional and genotype-dependent phenomenon, we calculated rostral/caudal ratios for densities estimated from our raw data (**Fig. 6E**). Plotting of the ratios confirmed that fractions for myelinating oligodendrocytes, OPCs and fibrous astrocytes were highly depleted in the rostral *Emx:MADM:F/F* areas, but appear to reemerge in the caudal SEZ/WM (**Fig. 6E**, values nearing 0). The absence of significant differences in ratios in the *Nes:MADM:F/F* SEZ/WM matches the more robust reemergence of green and red glia in these mice. Taken together these results support the possibility that dorsal deletion of *Egfr* (*Emx:MADM*) potently impacts rostrodorsal gliogenesis, whereas inclusion of ventral areas in the sparse recombination (*Nes:MADM* mice) reveals the reemergence of glia from a caudoventral domain in the forebrain.

Based on the robust and repeatable distribution patterns of MADM-labeled glia along the rostrocaudal axis of the cortex **(Fig. 7A)**, we next generated a heatmap with a dendrogram from a hierarchical clustering analysis to determine how cell densities compared in the regions analyzed for each genotype (**Fig. 7B**). We used densities of MADM-labeled neurons, glia (pooled oligodendrocytes and astrocytes), and progenitors broken down by region (rostral, caudal) and layer (UL, DL, SEZ/WM) for all six genotypes (*Emx:MADM* and *Nes:MADM*, +/+, *F/*+ and *F/F*). Clustering for cell types revealed that neurogenesis followed broadly similar patterns among genotypes and similarity was more influenced by region and layer than cell genotype, further suggesting that neurogenesis is largely an EGFR-independent process (**Fig. 7B**). In contrast, glial cell clustering was dramatically affected in *Nes:MADM:F/*+ and *Emx1:MADM:F/*+ cortices by cell genotype. Here, red glia formed a separate cluster from green glia, which correlated with the robust expansion of red glia and lack of green glia. A similar pattern was also apparent for cells within the SEZ/WM (S/W, **Fig. 7B**). These results indicate a surprising requirement for EGFR specifically in rostrodorsal gliogenic lineages that is apparently compensated for, at least in part, by a caudal domain when ventral progenitors are included in the MADM recombination in the *Egfr-null* background (i.e., in *Nes:MADM* cortices).

**Figure 7.**
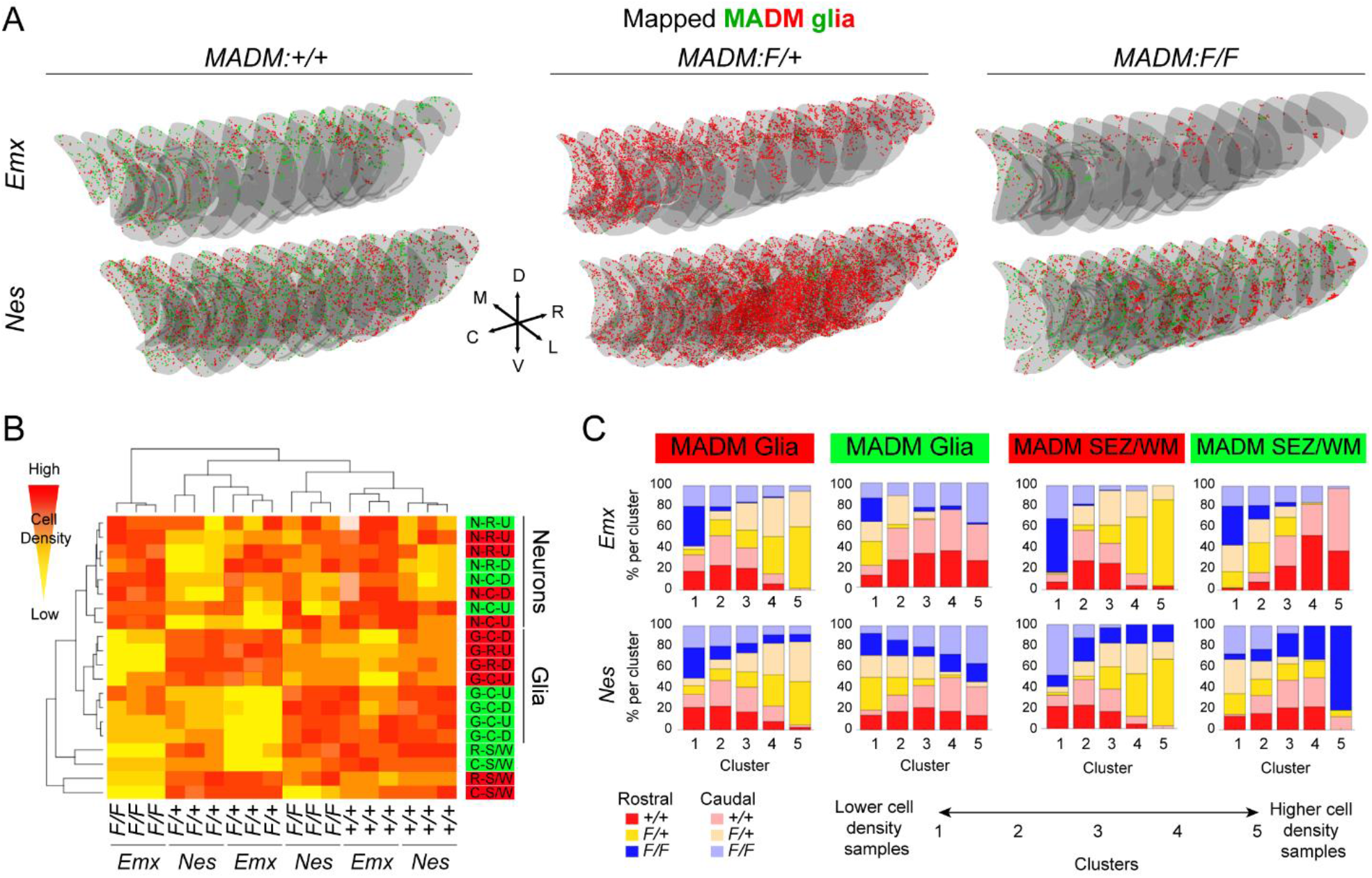
Distinct regional and laminar responses of gliogenic lineages *MADM:Egfr* cortices. **(A)** Reconstructions of red and green MADM glia in coronal sections through forebrains of P30 mice with the indicated genotypes. **(B)** Hierarchical clustering dendrogram of cell density measurements across MADM genotypes. Cell density measurements were used to generate heatmaps with hierarchical clustering. Red-to-yellow signals correlate with high-to-low relative cell densities. Clustering revealed neurogenesis is largely unaffected by *Egfr* dosage while glial and progenitor cell density is largely dictated by EGFR expression. Legend on the right of chart, color of highlight corresponds to MADM labeling (red or green); N, Neuron; G, Glia; R, Rostral; C, Caudal; S/W, Subependymal Zone/White Matter; U, Upper layer; D, Deep layer. (**C**) Self-organizing map unsupervised learning algorithm (SOM) was used to sort samples into linear arrays of five clusters. Clusterings were performed on subsets of data samples as indicated (tdTomato, red and GFP, green populations of glia and SEZ cells in *Emx:MADM* and *Nes:MADM* cortices).

To computationally and unbiasedly confirm the patterns illuminated by the hierarchical analysis, we carried out a clustering analysis on MADM cell density measurements (self-organizing map, SOM). Since we failed to observe any appreciable differences in neuronal populations (**Fig. S13**), separate clusterings were performed for densities of UL, DL and SEZ/WM glia in *Emx:MADM* and *Nes:MADM* cortices (**Fig. 7C**). In each clustering, the SOM grouped the cell densities into five ordered clusters, such that the samples with lowest cell densities were placed in the leftmost cluster (cluster 1) and greatest cell densities in the rightmost cluster (cluster 5; **Fig. 7C**). Since the SOM uses cell density measurements of each sample without labels for genotype (+/+ vs. *F/*+ vs. *F/F*)or location (rostral vs. caudal), the genotype and location of each sample can then be examined post-computationally in each cluster and correlated with cell densities in an unbiased manner.

The proportion of +/+ samples in each cluster generally increased from cluster 1 to cluster 5 (low green cell density to high green cell density). SOM clusterings of red MADM glia in the UL and DL revealed that they dominated clusters 4 and 5 with the highest densities in both *Emx:MADM and Nes:MADM* cortices (**Fig. 7C**). In contrast, *Egfr-null* glia in *F/*+ and *F/F* cortices were largely in clusters 1 and 2 (with low glial densities) in both cre lines. In comparison, clusterings of red SEZ/WM cells revealed that the *F/*+ samples (*WT* glia in an *Egfr-heterozygous* background) in clusters 4 and 5 with high densities were predominantly from the rostral domains of both *Emx:MADM* and *Nes:MADM* cortices. This suggested that the expansion of red glia in *F/*+ samples likely occurs in the rostral SEZ/WM. Matching our earlier observation, clusterings of *Emx:MADM* and *Nes:MADM* glia revealed that a substantial portion of red and green glia in cluster 5 (highest densities) came from caudal *F/F* samples (**Fig. 7C**). Interestingly, red and green *Nes:MADM:F/F* SEZ/WM cells in cluster 5 were predominately from rostral or caudal domains. This pattern suggests a highly asymmetric expansion of MADM clones during the gliogenic period in the *Nestin-cre* line which captures the ventral forebrain domain as well as the dorsal domain also covered in *Emx:MADM* cortices. The asymmetry in red and green populations was profoundly different in *F/F* samples, suggesting EGFR-independent expansion of glia may largely occur in one MADM sibling but not the other. Taken together, the SOM clusterings demonstrated that *Nes:MADM* SEZ/WM cells expand robustly, particularly in the rostral domain, in response to the *F/*+ and *F/F* genotypes, indicating their potential independence from EGFR in this population. The *Emx:MADM* condition provided for the observation that rostrodorsal progenitors in the forebrain lack the ability to either transition or expand into the gliogenic period in the absence of EGFR.

As revealed by *Emx:MADM:F/F*, an EGFR-dependent gliogenic progenitor population appears to be specifically present in the dorsal (cortical) domain. To further characterize the regional requirement for EGFR in the dorsal domain, we investigated *Emx:MADM* cell densities along the mediolateral axis of cortical areas in all three *Egfr* floxed genotypes (**Fig. S14)**. While lateral regions lacked glia, there was some recovery in more medial regions. Interestingly, the medial and caudal areas with substantial presence of glia in *Emx:F/F* forebrains corresponded to cortical tissues within, overlying and connected to the hippocampal formation (Anterior Cingulate, ACA; Retrosplenial, RSP; Rhinal; Subiculum, SUB; CA fields; dentate gyrus, DG). Further quantitative characterization and hierarchical analysis confirmed significant presence of glia in these areas within *Emx:F/F* forebrains, with the Dentate Gyrus containing the highest density of glia (**Fig. S14**).

Altogether, our findings indicate that gliogenesis, unlike neurogenesis, is robustly influenced by EGFR expression in the developing forebrain, yet there is a unique ventrocaudal EGFR-independent population of gliogenic progenitors that appears to compensate for loss of a rostrodorsal EGFR-dependent gliogenesis in the mouse forebrain.

## Discussion

In the current study we conducted a detailed quantitative and computational analysis of cortical development in various conditional *Egfr-*deletion models revealing that EGFR has a region-specific role in gliogenesis in the mouse forebrain. The requirement for EGFR was previously reported in germline *Egfr-*deleted mice which exhibit gross regional necrosis in the early postnatal forebrain, although the majority of them fail to survive past early postnatal life ^26^. In our study, most conditional *Nes:F/F* mice exhibit normal longevity following nursing and recovery during the pre-weaning age. We also noticed that *Nes:F/F* forebrain progenitor populations during the early postnatal period are dramatically reduced while cell death increased significantly in the olfactory bulbs. In fact, the size of the olfactory bulb remains significantly smaller in *Nes:F/F* compared to *WT* mice for the duration of their life. Moreover, the temporal pattern of EGFR expression reported in our study is consistent with previous observation of age-dependent differences in expression of various ErbB receptor family members (EGFR corresponds to ErbB1) in the mouse forebrain ^22^, and cortical progenitor cells ^24,55,56^. In addition, EGFR expression is upregulated between E16 and the first postnatal week in the cerebral cortices which is correlated with progressive elevation of mitotic responsiveness of progenitors to TGFα between E16 and E20 ^57^. Our findings using in situ panels from the Allen Brain Atlas, antibody staining and analysis of a new genetic *Egfr* reporter mouse, confirm the spatiotemporal expression patterns for EGFR in the forebrain. We extend this knowledge through the analysis of EGFR’s coexpression with a number of known biomarkers during perinatal forebrain development, where we found substantial overlap with a unique OLIG2/SOX9 dual expressing population. This population is rapidly depleted during early postnatal periods, suggesting that EGFR’s role on both oligodendrocytes and astrocytes through these putative bipotential progenitors is transient and rapid. Of the potential ligands for EGFR during this developmental period, TGFα, NRG1 and NRG2 exhibit appreciable expression levels and patterns in the forebrain. These patterns in the forebrain are suggestive of distinct roles for EGFR ligands that may correlate with the regional variation of effects of *Egfr* deletion on gliogenesis discovered in the current study.

### EGFR is required for generation of oligodendrocytes and homeostasis in the rostral forebrain

Our findings match the past linkage of EGFR signaling to the oligodendrocyte lineage using a hypomorphic *Egfr* mouse (*wave2*) and overexpression of EGFR ^24,25,29,31^. Development of forebrain oligodendrocytes is thought to originate from both the ventral and dorsal telencephalic regions, and OPCs display robust migratory and expansion capacities in the perinatal cortex and white matter, where they myelinate differentiating neuronal processes ^10^. The oligodendrocyte lineage can be partially distinguished from neuronal and other glial populations by expression of distinct markers during various developmental stages, similar to the progression that is seen in neuronal lineages. These include OLIG1 and OLIG2, PDGFRα, CC1 and NG2. While OLIG2 is necessary for specification of both oligodendrocytes and astrocytes during postnatal stages of development ^49,58^, OLIG1 is required for generation of myelinating oligodendrocytes ^59^. In addition, NG2^+^ cells contain a population of progenitors that give rise to both oligodendrocytes and a subset of protoplasmic astrocytes in the forebrain ^60^. Our finding that EGFR largely overlaps in its expression with OLIG1 and OLIG2 populations, but far less with PDGFRα and NG2 suggests that either EGFR/OLIG2/OLIG1 co-expressing cells are in a different stage of their development relative to PDGFRα and NG2 positive progenitors, or that they represent an independent population of gliogenic progenitors. The fact that both PDGFRα and NG2 cells are mildly affected in *Nes:F/F* cortices is suggestive of the latter possibility.

Further evidence for EGFR expressing cells constituting a unique population of gliogenic progenitors stems from the region-specific requirement for EGFR revealed in our study via the combination of bulk-deletion and MADM experiments. The potent requirement for EGFR in gliogenesis is associated with perinatal structural integrity of the rostrodorsal forebrain, especially in rostral neocortical areas, and the olfactory bulbs. Aligned with this finding, a past study illustrated that neuronal survival in the forebrain depends on EGFR signaling in cortical, but not midbrain astrocytes in cultures obtained from newborn *Egfr-null* mice ^33^. Remarkably, gross defects in these regions appear repaired during the first three weeks of postnatal life in *Nes:F/F* mice. However, a smaller size of the olfactory bulbs in 100%, and frontal cortical scarring in approximately 20% of *Nes:F/F* mice persist during their life-span. The main driving force behind these defects reveals itself in our comparison between *Nes:F/F* forebrains which contain both ventral and dorsal telencephalic deletion of *Egfr*, as opposed to *Emx:F/F* mice which only consist of dorsal deletion of *Egfr.* The most consistent defect in the *Nes:F/F* forebrain is the thinning of the white matter (i.e., the corpus callosum) concomitant with delayed emergence of myelin and the reduced thickness of myelin positive processes in these mice. The physiological effects of this defect on behavioral development and homeostasis in *Nes:F/F* mice remain to be elucidated.

### EGFR is required for protoplasmic astrocyte production in neocortical areas in a dosage-sensitive manner, whereas reactive astrocytes are EGFR-independent

Previous studies illustrated that enhanced expression of EGFR in the embryonic VZ leads to precocious emergence of astrocytes ^24^, and constitutive deletion of *Egfr* causes defects in astrocyte differentiation after birth ^27^. In our current study, effects on astrocyte production and maintenance appear genetically less penetrant in *Nes:F/F* forebrains than the effects on oligodendrocytes. Astrocytes perform diverse functions to support various brain processes through regulation of structural sustenance, water and ion balance, and the blood brain barrier ^61^. Astrocytes are classified into several groups that have recently become complex using single cell RNA sequencing results ^4,62–64^, but broadly they consist of fibrous, reactive, and protoplasmic astrocytes. The development process for these subtypes has not yet been demonstrated to follow a stepwise lineage as has been observed for neurogenesis and oligodendrocyte production, making it difficult to determine intermediate steps in the astrocyte differentiation process. Fibrous astrocytes display long unbranched cellular processes in the white matter and are molecularly distinguished from other glia by their high expression of GFAP. Astrocytes found in the grey matter are largely protoplasmic based on their supernumerary short processes and express significantly lower levels of GFAP. However, they are readily labeled with antibodies against the calcium-binding protein S100 and the transcription factor SOX9, and unlike reactive astrocytes, they establish anatomical domains that avoid neighboring astrocytes ^65^. At least a subset of GFAP positive astrocytes are ‘reactive’ to injury and disease by forming a scar to prevent further damage ^66,67^, though the source and diversity of ‘reactive glia’ remains unclear ^68^. For example, most reactive astrocyte responses are thought to occur locally ^69,70^ and are restricted to the lineage domains of origin (e.g., ventral versus dorsal) ^19^. However, there is also evidence that the ventral SVZ provide new reactive astrocytes into the cortical scar tissue ^71,72^. Interestingly, a small fraction of GFAP-positive astrocytes located in the SEZ/WM and another set in the adult hippocampus appear to function as neural stem cells in rodents and are essential for neuronal and glial production during adulthood ^73,74^. The degree to which EGFR expression impacts the subsets of astrocytes remains to be determined in future studies.

Our findings in *Nes:F/F* mice indicated that EGFR is not required for emergence and sustenance of GFAP-positive astrocytes in the SEZ/WM layers of the rostral or caudal cortices. In addition, the GFAP expressing population of reactive astrocytes/glia that responds to injuries also appear EGFR-independent in both young and aged mice, as stab injuries in the *Nes:F/F* rostral cortices results in GFAP-positive scarring similar to controls. Moreover, tissue necrosis accompanying the damage noted in the frontal cortices and the olfactory bulbs of *Nes:F/F* mice is consistently accompanied with clear GFAP-positive astrogliosis. As for protoplasmic astrocyte production in the *Nes:F/F* forebrain, a substantially lower number of SOX9 expressing astrocytes are present in the upper and lower layers of neocortical areas at P0. However, these numbers normalize to *WT* levels by P7 and are significantly higher by P30, corresponding to the scarring observed in the rostral forebrains of *Nes:F/F* mice. Together these findings indicate that the production and responsiveness of reactive astrocytes, fibrous astrocytes in the SEZ/WM, and protoplasmic astrocytes in the gray matter of the cortex are EGFR-independent. Whether the delayed emergence of protoplasmic astrocytes was a function of delayed maturation/migration of glioblast, or if an EGFR-independent population of gliogenic progenitors compensates for the loss of EGFR-dependent glia only became clear with the use of MADM.

### MADM reveals region-specific requirement for EGFR-dependent gliogenesis in rostrodorsal neocortical areas of the forebrain, and a novel EGFR-independent gliogenic population that can compensate for loss of EGFR

If EGFR is required for gliogenesis in the rostrodorsal domain of the telencephalon, how is it that *Emx:F/F* forebrains fail to exhibit any obvious phenotype? The answer to this became only possible through the use of MADM alleles and detailed quantification and analysis of GFP (green) and tdTomato (red) MADM sibling progenies in *Nes:MADM* and *Emx:MADM* cortices with the aid of two independent computational approaches. The results expose a novel EGFR-dependent gliogenic pattern that falls along the rostrocaudal and mediolateral extents of the cortex at P30, matching the *Nes:F/F* phenotypes resulting from bulk deletion of *Egfr* in these mice. We find a near complete abolition of glial-like cells in the *Egfr-null* (green) population in rostral cortical areas of both *Nes:MADM:F/*+ and *Emx:MADM:F/*+ mice. In contrast, there is a significant increase in both number and percentage of wildtype (red) glial populations in the same cortices (**Fig. 8; Movie S7**). These findings suggest that when faced with a largely *Egfr-heterozygous* forebrain background, sparse populations of *WT* cells overexpand to generate a larger number of glia than their normal capacity, whereas the *Egfr-null* MADM cells fail to generate glia in the same background. This overexpansion appears to be at least in part at the expense of neurons in the same clones (red MADM populations) in the *F/*+ background as there is a small but significant decline in neuronal populations in both *Nes:MADM:F/*+ and *Emx:MADM:F/*+ cortices. We found similar results in our recent clonal study using tamoxifen-inducible *NescreER:MADM:F/*+ alleles to label gliogenic progeny at clonal densities, and found a robust dosage response to EGFR expression in the generation of both astrocytes and oligodendrocytes ^18^. Since the red MADM glial progenitors are essentially overexpressing EGFR relative to their surrounding cells in the *MADM:F/*+ background, these results resemble gain-of-function findings that explain the strong effect of EGFR dosage on both astrocyte and oligodendrocyte production. Our findings further suggest a cell autonomous and EGFR-dependent role on astrocyte production that is likely compensated for by an EGFR-independent population in the bulk model described in the current study. Alternatively, glial progenitors may compete with one another such that the developmental depletion of one population (e.g., green *MADM:F*+ glia in our study) triggers proliferation in the other (e.g., red *MADM:F/*+glia). This process may not be controlled by EGFR levels, but rather by competition mechanisms that remain to be determined.

**Figure 8.**
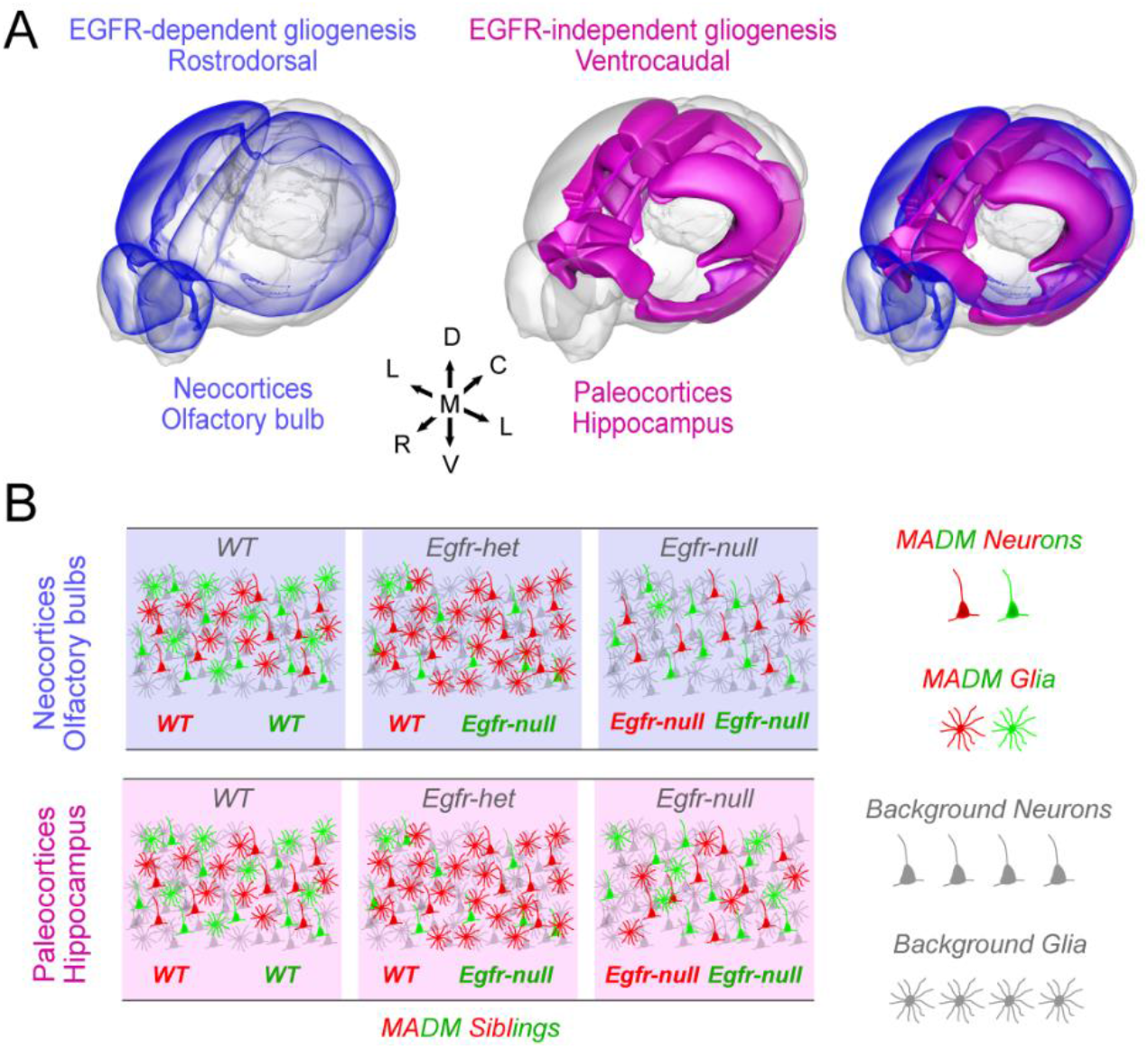
Summary. **(A)**Regional variation of EGFR dependency of gliogenesis along the rostrodorsal domains of the forebrain including the neocortices and the olfactory bulbs (blue). EGFR-independent gliogenesis depends on a novel ventrocaudal progenitor domain in the forebrain that normally supplies the paleocortical and hippocampal regions, but also appears to compensate for loss of EGFR-dependent gliogenesis in *Nes:MADM:F/F* and *Emx:MADM:F/F* forebrains described in the current study. **(B)** MADM analysis revealed a dosage dependency of regional gliogenesis on *Egfr*, and a novel EGFR-independent gliogenic population in the forebrain that supplies the paleocortical regions of the forebrain and appears to have the capacity to compensate for genetic loss of the EGFR-dependent population.

Another interesting and unexpected finding that stemmed from the *MADM:F/*+ analyses in the SEZ/WM was a robust elevation of fibrous astrocytes beyond a compensatory level (nearly double in number relative to controls; see **Figs. 6 and S12**). The lack of a similar effect on oligodendrogenesis in the SEZ/WM may suggest an absence of overdosage effect on this lineage which distinguishes them from SEZ/WM astrocytes. Another possibility is that the population of oligodendrocyte lineage may be highly dynamic and thus undergo stochastic expansion and elimination stages during the first four week of life in mice. In fact there is evidence for populations of OPCs that are developmentally transient and disappear in the adult mouse brain ^10,75,76^. Analyses of *MADM:F/*+ mice at earlier time points may shed further light onto these possibilities.

The most unexpected result in our current study emerges from the *Nes:MADM:F/F* forebrains where all the cells are presumably *Egfr-null.* Surprisingly, there is a reemergence of red and green MADM glia throughout the cortex, indicating they are derived from mitotic divisions rather than post-mitotic recombination events and hence developmentally derived. Although their densities are significantly lower in the frontal-most aspects of neocortical areas, red and green MADM glia are present in nearly all dorsal and ventral territories of the forebrain including neocortical and paleocortical areas. Glia are also found in *Emx:MADM:F/F* forebrains but largely in the hippocampal formation and some of its associated cortices in the medial and caudal aspects of the dorsal telencephalon. The comparison of this finding needs to be placed in the context of current knowledge on the regional origin of astrocytes. A previous study showed that dorsal and ventral progenitors in the forebrain participate in astrocyte production in a restricted manner ^19^. Accordingly, dorsal progenitors that largely give rise to excitatory projection neurons in the forebrain only produce dorsal astrocytes (largely in the cerebral cortices and hippocampus), whereas ventral progenitors which produce both ventral and dorsally seeded neurons (in subcortical forebrain nuclei and inhibitory neurons in the cortex) only produce ventrally seeded astrocytes.

While astrocytes derived from ventral and dorsal progenitors in the telencephalon maintain their positional identity by maturity ^9^, positional identity of oligodendrocytes in the cortex is more complex and may involve multiple ventral and dorsal progenitor domains ^10^. In this context, the comparison of our *Emx:MADM:F/F* and *Nes:MADM:F/F* results raises the possibility that an EGFR-independent gliogenic progenitor pool, likely positioned in the ventrocaudal aspect of the forebrain, appears capable of compensating for the loss of EGFR-dependent glia. Surprisingly, the EGFR-dependent glia appear to occupy neocortical areas, whereas the EGFR-independent populations largely revealed in comparison of *Emx:MADM:F/F* to *Nes:MADM:F/F* cortices, appear to seed evolutionary older paleocortical areas including the hippocampal formation along the rostromedial and caudal aspects of the forebrain (**Fig. 8; Movie S7**). Interestingly, we failed to observe any crossing of the dorsal *Egfr-null* glia into ventral forebrain territories of *Emx:MADM* mice, whereas the *Nes:MADM:F/F* result raises the possibility of substantial crossing of ventral gliogenic progenitors or glioblast into the dorsal telencephalon during perinatal development in mice. This raises the possibility that an EGFR-dependent population of glia likely participates in production of both ventral and dorsal glia. It appears that an EGFR-independent gliogenic population in the ventrocaudal aspect of the *Nes:F/F* forebrain attempts to compensate for loss of the EGFR-dependent progenitors that populate the rostrodorsal domains of the forebrain (**Fig. 8; Movie S7**). The precise identity and positioning of the EGFR-independent glia producing progenitors remains to be determined.

Our study raises a number of important questions. What are the inducers and regulators of the sudden and potent upregulation of EGFR during the gliogenic switch in the forebrain? How are the oligodendrocyte and astrocyte lineages linked in different forebrain regions such as the paleocortical and neocortical boundaries? What is the developmental source and molecular identity of the putative EGFR-independent glial populations identified in our study? Are they related to the reactive glial populations that persist throughout life? Do ventral gliogenic populations migrate into the cortex to compensate for loss of the EGFR-dependent gliogenic progenitors in *Nes:F/F* cortices, and if so, when and how? We have begun to address some of these questions by implementing automated and unbiased technologies into analysis of the forebrain ^77^. Future studies addressing these questions will enhance our understanding of the important developmental process of gliogenesis and its regulation of forebrain homeostasis and function. Finally, *Nes:F/F* mice utilized in our study are able to live up to two years with early postnatal care, and when maintained under homeostatic laboratory housing conditions providing a unique mouse model to study glial functions and dysfunction in the absence of EGFR signaling. Consequences of perinatal defects, the nature of perinatal repair and possibly regeneration in the olfactory bulbs and the frontal cortices, and lack of sufficient oligodendrocyte production on behavioral paradigms will be intriguing next steps in characterization of this unique mouse line.

### Limitations of the Study

A limitation of the current findings stems from our incomplete understanding of the cre lines used in our study and the degree to which MADM captures all progenitor populations in the developing forebrain. For example, it is possible that neither Nestin nor Emx1 cre lines encompass the entirety of ventral and dorsal progenitor populations in the telencephalon. As such, it is possible that we fail to identify the role of EGFR in those missed populations. Secondly, the rare mitotic recombination events (e.g., x- and z-segregations) underlying MADM may only occur in subpopulations of progenitors, again possibly missing the role of *Egfr* in unlabeled cells. While we think these are unlikely based on the robust and highly repeatable phenotypes in our models, future genetic approaches may be considered to address these possibilities.

## Acknowledgments

The authors are grateful to Drs. David Rowitch (UCSF/Cambridge) for the OLIG1, Dwight Bergles (Johns Hopkins) for the NG2, and Benjamin Deneen (Baylor College of Medicine) for the NFIA antibodies. The authors thank members of the Ghashghaei lab for discussions, and Dr. Mahdi Aliomrani for careful review of the manuscript.

## Author contributions

Conceptualization, H.T.G.; Methodology, H.T.G, X.Z., G.X., D.T., R.C., A.G., R.C., X.P., R.E., and M.C.O.; Investigation, H.T.G., X.Z., G.X., C.A.J., and Z.K.H.; Visualization, H.T.G., G.X., A.G., and R.C.; Supervision, H.T.G.; Writing – Original Draft: H.T.G. and G.X.; Writing – Review & Editing, H.T.G., G.X., C.A.J., M.C.O., A.G., D.T., and Z.K.H.

## Data and materials availability

All data needed to evaluate the conclusions in the paper are present in the paper and/or the Supplementary Materials. Additional raw data, code, and materials used in the analyses will be made available to any researcher for purposes of reproducing or extending the analyses. *Egfr^F/F^* and *Egfr^EM^* mice should be requested from DT and RC, respectively.

---

**Movie S1. Related to Figure 5.**

A 3D reconstruction of an intact tissue cleared hemisphere of an *Emx:MADM:*+/+ mouse forebrain. The movie shows consecutive sagittal digital sections (30 μm thickness) at different depths. The hemisphere was imaged using a custom-built light sheet fluorescence microscope.

**Movie S2. Related to Figure 5.**

A 3D reconstruction of an intact tissue cleared hemisphere of an *Emx:MADM:F/*+ mouse forebrain. The movie shows consecutive sagittal digital sections (30 μm thickness) at different depths. The hemisphere was imaged using a custom-built light sheet fluorescence microscope.

**Movie S3. Related to Figure 5.**

A 3D reconstruction of an intact tissue cleared hemisphere of an *Emx:MADM:F/F* mouse forebrain. The movie shows consecutive sagittal digital sections (30 μm thickness) at different depths. The hemisphere was imaged using a custom-built light sheet fluorescence microscope.

**Movie S4. Related to Figure 5.**

A 3D reconstruction of an intact tissue cleared hemisphere of an *Nes:MADM:*+/+ mouse forebrain. The movie shows consecutive sagittal digital sections (30 μm thickness) at different depths. The hemisphere was imaged using a custom-built light sheet fluorescence microscope.

**Movie S5. Related to Figure 5.**

A 3D reconstruction of an intact tissue cleared hemisphere of an *Nes:MADM:F/*+ mouse forebrain. The movie shows consecutive sagittal digital sections (30 μm thickness) at different depths. The hemisphere was imaged using a custom-built light sheet fluorescence microscope.

**Movie S6. Related to Figure 5.**

A 3D reconstruction of an intact tissue cleared hemisphere of an *Nes:MADM:F/F* mouse forebrain. The movie shows consecutive sagittal digital sections (30 μm thickness) at different depths. The hemisphere was imaged using a custom-built light sheet fluorescence microscope.

**Movie S7. Related to Figure 8.**

Video of a rotating 3D cartoon of a mouse forebrain delineating neocortical areas (blue) where gliogenesis is EGFR-dependent, and paleo regions (purple) including the hippocampal forebrain in which gliogenesis appears EGFR-independent. Cartoon was generated in the Scalable Brain Alas, using the 3D mouse brain module (https://scalablebrainatlas.incf.org/composer/?template=ABA_v3) defined with our quantitated MADM data.

## STAR★ Methods

### Resource Availability

#### Lead Contact

Further information and requests for resources and reagents should be directed to and will be fulfilled by the lead contact, H. Troy Ghashghaei.

#### Materials Availability

- This study did not generate new unique reagents.

#### Data and code availability

- RNA sequencing data have been deposited at Gene Expression Omnibus and are publicly available as of the date of publication. Accession numbers are listed in the key resources table.
- All data reported in this paper will be shared by the lead contact upon request.
- This paper does not report original code.
- Any additional information required to reanalyze the data reported in this paper is available from the lead contact upon request.

### Experimental Model and Subject Details

Mice were used under the regulations and approval from Institutional Animal Care and Use Committee at North Carolina State University. Mice were housed in a 12-hour light:dark cycle with ad libitum access to food and water. To delete *Egfr* in the brain, we utilized *Nestin-Cre* (https://www.jax.org/strain/003771 or *Emx1^cre^* (https://www.jax.org/strain/005628) mice (both on a mixed C57BL/6/SJL backgrounds) crossed to *Egfr floxed* mice (*Egfr^F/F^*)^78^. Cre-mediated recombination in neural progenitors results in deletion of *exon 3* of *Egfr* causing a frame shift that generates two stop codons terminating transcription. The following genotypes were used for bulk-deletion analyses: *Wildtype* (*WT*: *Egfr^+/+^* or *Nestin-Cre:Egfr^+/+^*), conditional heterozygous for *Egfr* deletion (*Nes:F/F Nestin-Cre:Egfr^+/+^*), and conditional homozygous for *Egfr* deletion (*Nes:F/F*); *Nestin-Cre:Egfr^F/F^*). Mosaic Analysis with Double Marker (MADM) mice were generated by intercrossing of *MADM11^TG^* and *MADM11^GT^* mice (https://www.jax.org/strain/022977, and https://www.jax.org/strain/022976). Final MADM mice for imaging and analyses were obtained by crossing *Egfr^F/+^;MADM-11^TG/TG^* or *Egfr^F/F^;MADM-11^TG/TG^* mice to *Nestin-cre;MADM-11^GT/GT^ or Emx^cre^;MADM-11^GT/GT^*. Control mice were generated by crossing *Nestin-cre;MADM-11^GT/GT^ or Emx^cre^;MADM-11^GT/GT^ mice to MADM-11^TG/TG^* mice. *Egfr floxed* alleles were detected by PCR using the primers: *Egfr lox3* F: 5’ CTTTGGAGAACCTGCAGATC; *Egfr lox3* R: 5’ CTGCTACTGGCTCAAGTTTC. All other mouse lines (*MADM-11 alleles* and *Nestin-cre*) were genotyped using protocols provided by Jackson Laboratories.

### Method Details

#### BrdU Labeling, Histochemistry and Immunohistochemistry

5’-Bromo-2’-deoxyuridine (BrdU) is a thymidine analog that integrates into newly synthesized DNA in dividing progenitors upon administration *in vivo* and *in vitro*. Prior to perfusion, mice were intraperitoneally injected with BrdU dissolved in 0.9% saline at a concentration of 10 mg/ml and administered at 100 μg/g body weight. One hour later, mice were perfused with 4% Parafomaldehyde and the brains were removed from the skull and post-fixed overnight. Brains were sectioned into 50 μm sagittal sections with a vibratome. For immunohistochemistry, floating brain sections were blocked with 10% goat serum and 1% Triton-X 100 in 0.1 M PBS for 1 hour at room temperature. Sections were then incubated at 4°C overnight with one or a combination of antibodies listed in Key Resources Table. All primary antibodies were diluted in PBS with 1% goat serum, 0.3% Triton X-100. Sections were washed in PBS three times (five minutes each), followed by incubation with species-specific conjugated fluorescence secondary antibodies for one hour at room temperature. After secondary antibody incubation, the sections were washed with the same washing protocol and coverslipped using Fluoromount Aqueous Mounting Medium (MiliporeSigma).

Weil’s stain was utilized for histochemical labeling of myelin. Briefly, vibratomed sections were mounted on slides and hydrated using standard alcohol series (95%-75%-50%, four minutes each). Slides were then placed in distilled water for 10 minutes followed by rinse for 30 minutes in preheated (50-60°C) staining solution (Haemotoxylin diluted 1:10 in absolute ethanol with 4% ferric ammonium sulphate). Sections were then washed in tap water for 10 minutes followed by partial differentiation for five minutes in Weigert’s differentiator solution (1% Borax and 1.25% potassium ferricyanide). Sections were washed three times for five minutes each wash in distilled water followed by completion of differentiation in Wigert’s differentiator for two minutes. Following a final wash series in distilled water the sections were dehydrated in reverse alcohol series followed by clearance with Xylene and coverslipped with Permount (Fisher).

#### RNA sequencing and bioinformatics analyses

E17.5 embryonic and P5 postnatal *Nes:F/F* and control (*Egfr^F/+^* or *Nestin-cre*) mice were obtained from time-pregnant females. The day of vaginal plug detection was considered E0.5. Mice were collected in fresh ice cold HBSS and brains were taken out of the skull, and forebrains were separated from the rest of the brain using fine microforceps in ice cold HBSS. The cortices and the hippocampal formation were dissected away from the ventral forebrain using microforceps and split into rostral and caudal halves. Samples were collected and Total RNA was extracted using the RNeasy Mini kit (Qiagen, catalog no. 74104). All samples were quantified and assayed to confirm minimum RNA integrity number of at least 9.3 using an Agilent Bioanalyzer (High Sensitivity DNA Kit, catalog no. 5067-4626). Next, 1 μg of total RNA per sample underwent mRNA capture and was then fragmented at 94 °C for 6 min. Sequencing libraries were prepared according to the manufacturer’s protocol using ten cycles of final amplification (KAPA mRNA HyperPrep Kit, catalog no. KK8580 and KAPA UDI Adapter Kit, catalog no. KK8727). Next-generation sequencing was performed on an Illumina NextSeq500 (75-bp paired end) to a targeted depth of ~20 million reads per sample.

#### Tissue clearing and light sheet microscopy

Brain hemispheres were first dehydrated using methanol gradients (20%, 40%, 60%, 80%, 100% in dH_2_O) 1 hour each at RT, then washed with 100% methanol for 1 hour at RT and chilled for 2 hours at 4 °C. Next, chilled brain hemispheres were submerged in 66% dichloromethane/33% methanol at RT overnight. Hemispheres were then washed twice in 100% methanol at RT, chilled for 2 hours at 4 °C and bleached in chilled fresh 5% H_2_O_2_ in methanol at 4 °C overnight. After bleaching, brain hemispheres were rehydrated with methanol series (100%, 80%, 60%, 40%, 20% in dH_2_O) 1 hour each at RT and washed with PBS for 1 hour at RT, followed by two washes with PBS with 0.02% Triton X-100, 1 hour each at RT. Next, the hemispheres were permeabilized with 0.2% Triton X-100, 0.3M Glycine, 20% DMSO in PBS for 2 days at 37 °C, then incubated in blocking solution (0.2% Triton X-100, 6% donkey serum, 10% DMSO in PBS) for 2 days at 37 °C. Samples were then incubated with chicken anti-GFP (1:200, Aves Labs) and rabbit anti-RFP (1:200 Rockland) primary antibodies in PTwH (0.2% Tween-20, 10 mg/L heparin in PBS) with 5% DMSO and 3% donkey serum for 5 days at 37 °C. Hemispheres were washed 5 times with PTwH, 2 hours each until the next day. Brain hemispheres were then incubated with AlexaFluor donkey anti-chicken 647 (1:200, Jackson ImmunoResearch) and donkey anti-rabbit Cy3 (1:200, Jackson ImmunoResearch) in PTwH with 3% donkey serum for 5 days at 37 °C, followed by 5 washes with PTwH. For optical clearing, brain hemispheres were dehydrated with methanol series (20%, 40%, 60%, 80%, 100% in dH_2_O) 1 hour each and incubated in 66% dichloromethane/33% methanol for 3 hours, both at RT. Next, brain hemispheres were submerged in 100% dichloromethane 15 minutes twice at RT. All the above steps were performed with gentle shaking. Brain hemispheres were submerged in DiBenzyl Ether until they became fully transparent. Cleared brain hemispheres were imaged using a custom-built light sheet microscope, the setup of which was outlined in ^79,80^. Acquired images were stitched using TeraStitcher ^81^ and then visualized in Imaris 9.5 (Oxford Instruments) for preparing the movies through each cleared and stained hemisphere.

#### Neurosphere Cultures

Brains were collected rapidly from hypothermically anesthetized newborn *WT* and *Nes:F/F* mice. Individual cells were dissociated from wholemount preparations of the SEZ ^82^, followed by the enzymatic dissociation as described previously ^83^. Dissociated cells were cultured at one million cells per well in 6-well Fugene culture dishes, in 1.5 mL of Neurobasal medium supplemented with hEGF (1 ng/mL, Invitrogen) and bFGF (10 ng/mL, Invitrogen). The growth factors were added every two days and the obtained neurospheres were passaged by dissociation and re-culturing every 4-6 days.

#### Western Blotting

Forebrains from control (either *Egfr^F/F^* or *Nestin-cre* mice) and *Nes:F/F* mice were rapidly dissected in lysis buffer (50mM Tris HCl, pH 8.3; 1% Triton-X, 500mM EDTA, pH 8.0; 100mM NaCl, 50mM NaF) with protease inhibitor cocktail (Roche) and homogenized for 3 minutes with a bullet blender. The BCA assay was performed on total protein according to the manufacture’s protocol (Pierce BCA kit). Protein samples were run in polyacrylamide gel and transferred to nitrocellulose blots. The blots were processed for *Egfr* detection using standard Western blotting protocols. Blots were developed with the ECL kit (Pierce).

### Quantification and Statistical Analysis

Sections were imaged on a Nikon Eclipse EZ-C1 or Olympus FV1000 confocal microscopes. Immunolabeled cells in forebrain sections were quantified using standard stereological estimation methods as described previously ^84^. Statistical significance was determined using Student’s t-test and all values were expressed as mean ± standard error of mean (SEM). The grey scale images illustrating the prevalence of SOX9/OLIG2 double-labeled nuclei in Fig. S6 were obtained by thresholding, which was conducted in Adobe Photoshop using the Hue/Saturation function to eliminate the red and green signals, leaving only the yellow, double labeled, signal. The thresholded images were desaturated to yield the greyscale images.

Thickness of the Myelin positive corpus callosum or WM was measured in brightfield images captured using a 10x lens in regionally matched sections from *WT* and *Nes:F/F* forebrains. Measurements were conducted in ImageJ using the line measurement tool, spanning Myelin positive tissue from the ventricular surface extending basally. Data were transferred to Microsoft Excel for calculation of ratios and mean ratios, and corresponding SEM’s were plotted for presentation.

Signal intensities for GFAP immunostained brain sections were quantified in Image J using gray scale images in single slices of Z-stack confocal images captured using 10x or 20x objectives. Measurements were obtained within cortices 500 μm rostral and 500 μm caudal to the site of injury. Recordings were transferred to an Excel file for analysis. Background measurements on the same images from surrounding tissue with no fluorescent/chromatic labeling were subtracted from readouts. Signal intensities were presented on a normalized scale of 0 (no signal) to 250 (fully saturated signal).

Neurospheres were imaged on a Nikon EZ-C1 confocal microscope once a day throughout all passages and the number of neurospheres were counted in five random 2 mm^2^ grids in each culture well for analysis.

Power analysis (P=90; α=0.01) was conducted for analysis of MADM cells to determine ideal sample sizes across the two different cre lines (*Nestin* and *Emx*) and the three *Egfr* genotypes (*+/+, F/+, F/F*). Comparison of GFP and tdTomato positive MADM cells in both the *Nestin* and *Emx* cre lines on the *Egfr^F/+^* background with clear difference in red and green population of glia indicate that for the grid sizes used in our study (300 μm × 300 μm), we required a maximum of 104 samples to obtain enough MADM neurons in the upper and deep layers of the cortex. Repeating this analysis for glia indicated that a maximum of 94 sampling grids were required. We used a total of 105 sampling grids for each layer spread across three animals per genotype.

Counting of MADM cells was conducted in two ways in the layers of various cortical areas. First, all MADM cells were counted in five sagittal sections containing rostral and caudal forebrain regions per mouse per genotype containing intact olfactory bulbs. Second, samples in sagittal and coronal sections were counted using 300 μm x 300 μm grids randomly placed in different cortical areas until 105 total sampling grids were counted across three animals per genotype. Hierarchical heatmaps were generated in RStudio using the package gplots ^85^. Both packages were run using default parameters.

#### Computational Analysis of MADM Data

Clustering analysis was applied to MADM data using a self-organizing map (SOM), an artificial neural network (ANN) algorithm that is an extension of k-means clustering accounting for topology via neighborhood relationships between clusters. Specifically, a linear array of five clusters was employed: three interior clusters, each with two neighbors, and two end clusters, each with a single neighbor. Each cluster in the SOM was assigned a weight vector having the same number of components as the data to be clustered. Weights were tuned during the training phase by: (i) presenting random batches of data to the network; (ii) finding the closest (winning) cluster for each data vector and adjusting the weight vector for this cluster to move in the direction of the data vector; (iii) moving the weight vectors for neighboring cluster(s) in the direction of the data vector, but with a smaller amplitude applied to the differential vector. These steps were repeated as a learning rate parameter was systematically decreased, thus also reducing the amplitude of the weight vector adjustments during training. The result was a mapping of data vectors to five distinct clusters. Due to the linear arrangement (topology) of the SOM, the trained ANN effectively provides an unbiased sorting of the data from which trends can be identified by contrasting patterns clustering to opposite sides of the linear array of five clusters on the map. A similar approach using the same ANN architecture was applied for data sets in other applications ^86^.

#### Bioinformatics Analysis

For bioinformatics analyses, raw reads were adapter trimmed with BBDuk 38.94 ^87^ and aligned to the GENCODE GRCm39 primary assembly (release M29) using STAR 2.7.8a ^88^. Aligned reads were quantified with featureCounts 2.0.3 ^89^ and normalized to TPM. The resulting gene expression matrix was inspected for outliers with the SampleNetwork R function ^90^ and batch corrected using ComBat ^91^. Multiple linear regression was performed for all genes after removing genes below the 30th percentile of mean expression values (n=26,281). Expression values were scaled in the gene dimension and modeled by factors for genotype, region, and their interaction. Genes were ordered in ascending or descending order by regression coefficient for each model term and input to the GSEA function from the clusterProfiler R package (version 4.2.2) ^92^. For a gene set to be found significant with respect to a given covariate (genotype, region, or interaction) for a given direction of association (positive or negative), it had to be more significantly enriched for that covariate than the other two for the same direction of association. To identify modules of highly correlated genes, coexpression analysis was performed as described ^93^. Briefly, pairwise biweight midcorrelations were calculated on the batch-corrected gene expression matrix. Genes were clustered using a hierarchical clustering procedure with complete linkage and (1–bicor) as a distance measure. The resulting dendrogram was cut at a series of heights, enforcing varying minimum module size to requirements, yielding a total 35 different gene coexpression networks. Within each network, similar modules were merged if the correlations of their module eigengenes (PC1) exceeded an arbitrary threshold (0.85). This procedure was performed iteratively for each network, until no pairs of modules exceeded the threshold. The resulting modules were then characterized using a one-sided Fisher’s exact test to test for significant gene set enrichments.

## KEY RESOURCES TABLE

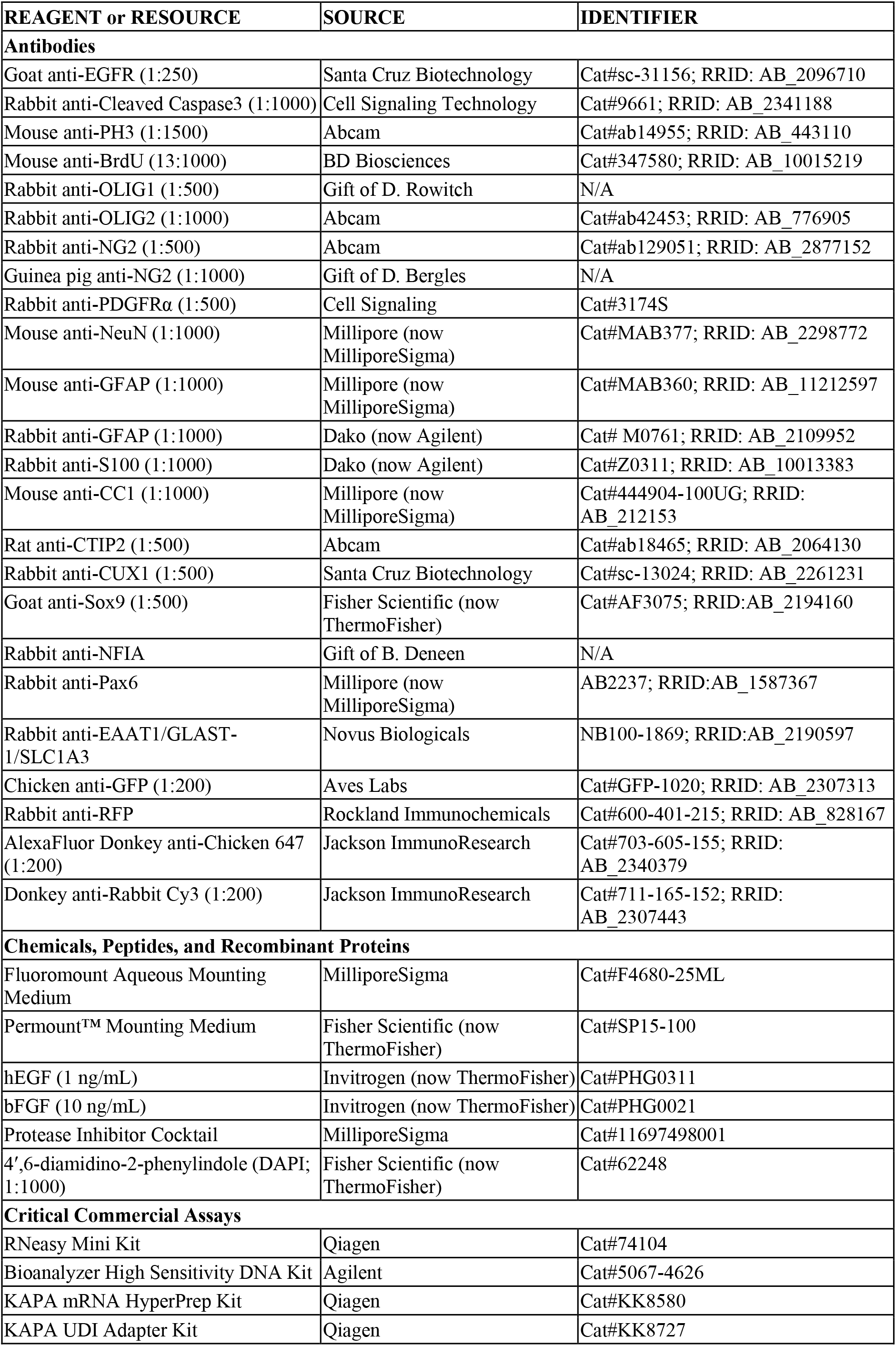
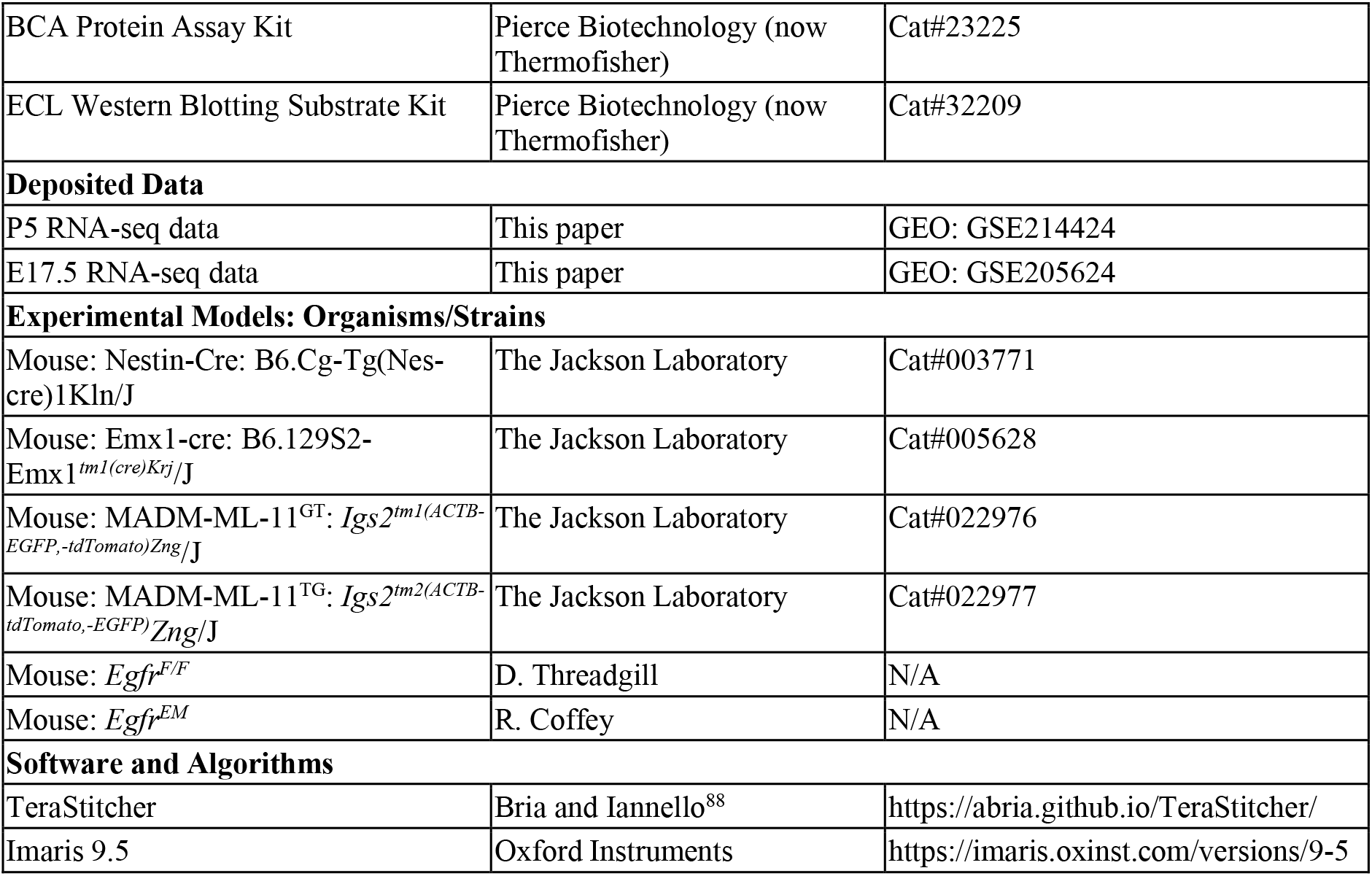

