## Supplemental figures for "Bulk and Mosaic Deletions of *Egfr* Reveal Regionally Defined Gliogenesis in the Developing Mouse Forebrain"

### Title

**Short title:** Regional role of *Egfr* for Gliogenesis in the Forebrain


### Authors

Xuying Zhang [1,7], Guanxi Xiao [1,7], Caroline Johnson [1], Yuheng Cai [2], Zachary K. Horowitz[1], Christine Mennicke [3], Robert Coffey [4], Mansoor Haider [4], David Threadgill [5], Rebecca Eliscu [6], Michael C. Oldham [6], Alon Greenbaum [2], H. Troy Ghashghaei [1,8]

### Affiliations

1. Department of Molecular Biomedical Sciences, North Carolina State University, Raleigh, NC, USA
2. Joint Department of Biomedical Engineering, North Carolina State University and University of North Carolina at Chapel Hill, Raleigh, NC, USA
3. Department of Mathematics, North Carolina State University, Raleigh, NC, USA
4. Department of Medicine, Vanderbilt University Medical Center, Nashville, TN, USA
5. Institute for Genome Sciences and Society, Texas A&M University, College Station, TX, USA
6. Department of Neurological Surgery, University of California at San Francisco, San Francisco, CA, USA

7 Equal Contribution



### Summary

The Epidermal growth factor receptor (EGFR) plays a role in cell proliferation and differentiation during healthy development and tumor growth, however its requirement for brain development remains unclear. Here we used a conditional mouse allele for *Egfr* to examine its contributions to perinatal forebrain development at the tissue level. Subtractive bulk ventral and dorsal forebrain deletions of *Egfr* uncovered significant and permanent decreases in oligodendrogenesis and myelination in the cortex and corpus callosum. Additionally, an increase in astrogenesis or reactive astrocytes in effected regions was evident in response to cortical scarring. Sparse deletion using Mosaic Analysis with Double Markers (MADM) surprisingly revealed a regional requirement for EGFR in rostrodorsal, but not ventrocaudal glial lineages including both astrocytes and oligodendrocytes. The EGFR-independent ventral glial progenitors may compensate for the missing EGFR-dependent dorsal glia in the bulk *Egfr*-deleted forebrain, potentially exposing a regenerative population of gliogenic progenitors in the mouse forebrain.

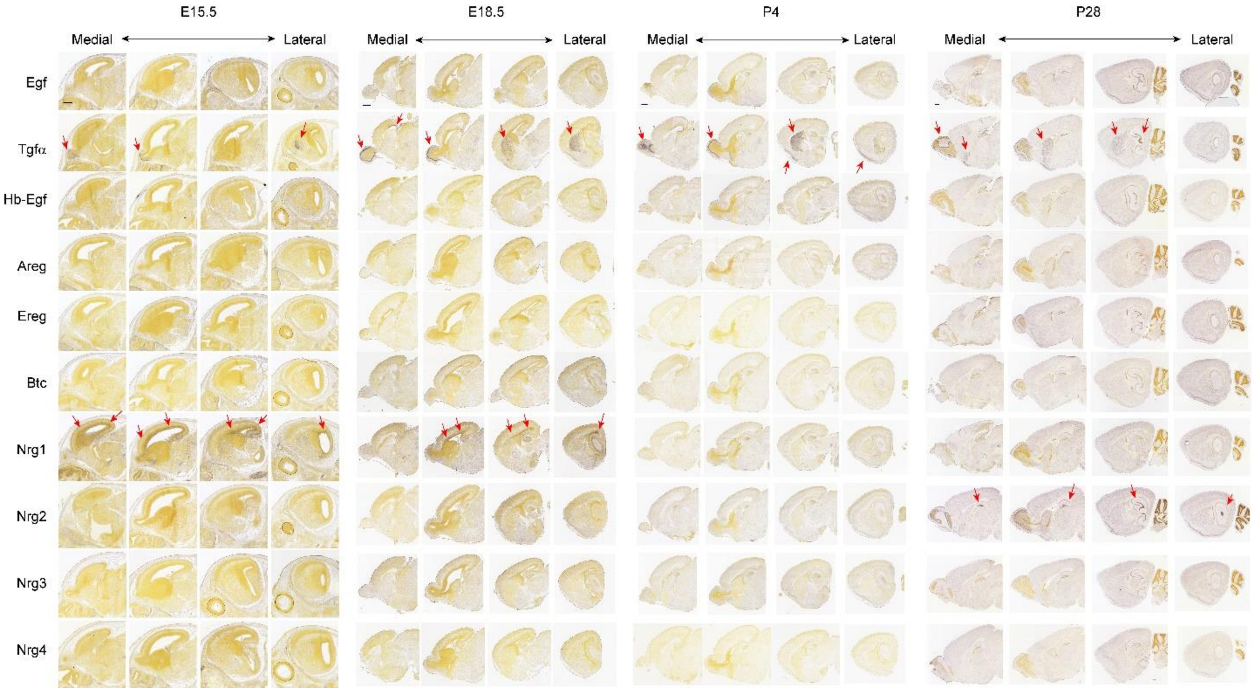


**Figure S1. In situ hybridization panels for known EGFR ligands from the Allen Brain Atlas. Related to Figure 1. Expression of known EGFR ligands.** Egf, Epidermal growth factor; Tgfα, Transforming growth factor α, Hb-Egf, Heparin-binding Egf; Areg, Amphiregulin; Ereg, Epiregulin; Btc, Betacellulin; NRG1-4; Neuregulin 1-4. Red arrows point to appreciable region-specific signals in the forebrain for Tgfα, Nrg1, and Nrg2. Scale bars, 500 µm for each age.

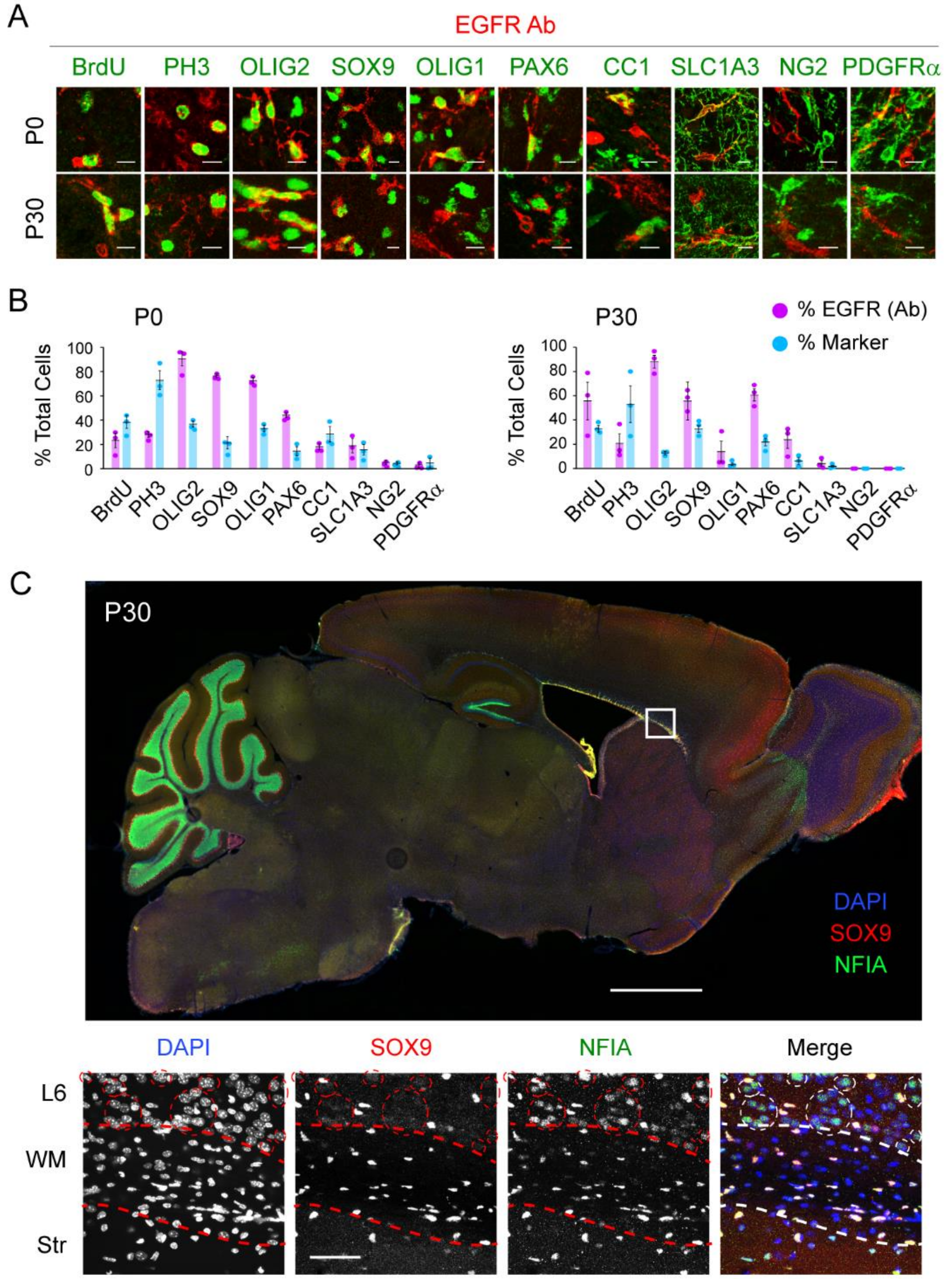


**Figure S2. Characterization of EGFR and NFIA antibody staining in the early postnatal forebrain. Related to Figure 1. (A)** Overlap of EGFR antibody-stained cells with various markers in the SEZ/WM of mice at indicated stages. (**B**) Bar charts present proportions of cells positive for each marker as percentages of total EGFR-positive or marker-positive populations as indicated. Scale bar, 50 µm. (**C**) Low power image of a sagittal section co-immunostained for Sox9 (red) and NFIA (green) and counterstained for DAPI (blue). High power images below is from boxed area in the low power image. The three colors are provided as desaturated images for each marker and merged as indicated. Tick dotted line demarcates the striatum (Str), WM, and layer 6 (L6) of the cortex. Dotted circles indicate NFIA positive neurons in L6.

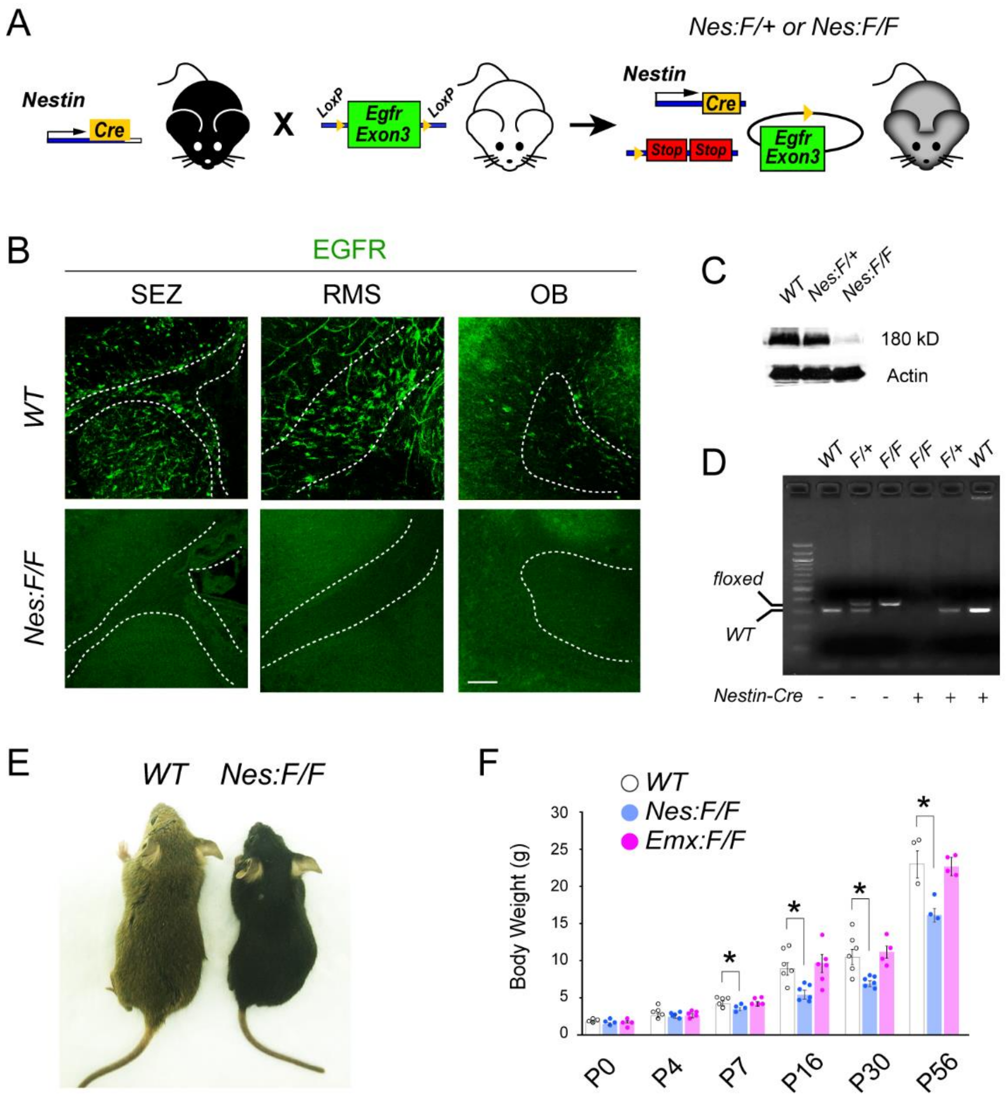


**Figure S3. Assessment of EGFR expression in the *Nes:F/F* forebrain. Related to Figures 2-3. (A)** Conditional alleles for Egfr (*exon 3* flanked by *loxP* sites) were crossed to *Nestin-cre* transgenic mice to generate *Nes:F/F* animals. **(B)** Confocal images of Egfr antibody-stained tissue in *WT* and *Nes:F/F* forebrains at P7. Note the absence of EGFR immunoreactivity in the SEZ, RMS, and OB of *Nes:F/F* forebrains. Dotted lines delineate the boundaries of each structure. Scale bar, 50 µm. **(C)** Western blotting against EGFR in lysates obtained from *WT*, *Nes:F/+*, and *Nes:F/F* forebrains at P7. **(D)** PCR results on *Nestin-cre* positive and negative forebrain samples subject to PCR detection of *floxed* and *WT* alleles for *Egfr* across the three floxed mouse genotypes (*WT, F/+, F/F*). Left lane, 100 bp ladder. **(E)** A typical pair of *WT* and *Nes:F/F* littermates at P21. **(F)** Body weights of *WT*, *Nes:F/F*, and *Emx:F/F* mice across various ages. Data are mean ± s.e.m; individual dots are for each n. Asterisks, Student's t-test, $p < 0.05$.

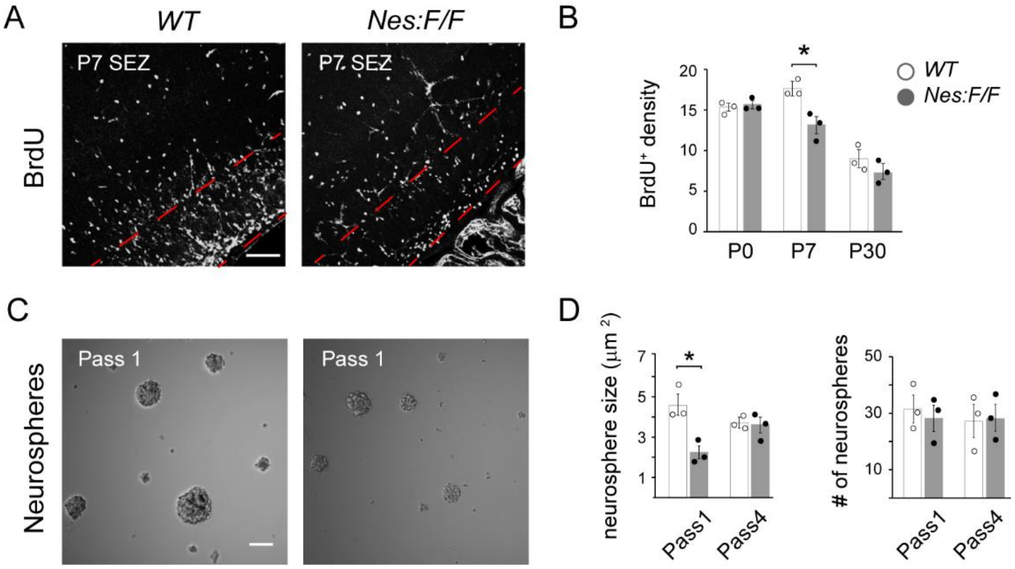


**Figure S4. Progenitor proliferation in the *Nes:F/F* forebrain during early postnatal development. Related to Figures 2-3. (A)** Confocal micrograph of BrdU labeled tissue around the SEZ at P7. Scale bar, 50 µm. **(B)** Densities of BrdU+ cells in *WT* control and *Nes:F/F* SEZs at P0, P7, and P30. Data are mean ± s.e.m of densities (x $10^5$ / $mm^3$), n=3 animals. **(C)** Representative micrographs of neurospheres grown from SEZ/RMS extracts after the first passage in the presence of EGF and FGF. Scale bar, 100 µm. **(D)** Average size and number of neurospheres between *WT* and *Nes:F/F*. Data are mean ± s.e.m of neurospheres size and number of neurospheres per counting area ($10^5$ $µm^2$). *, Student's t-test, $p < 0.05$.

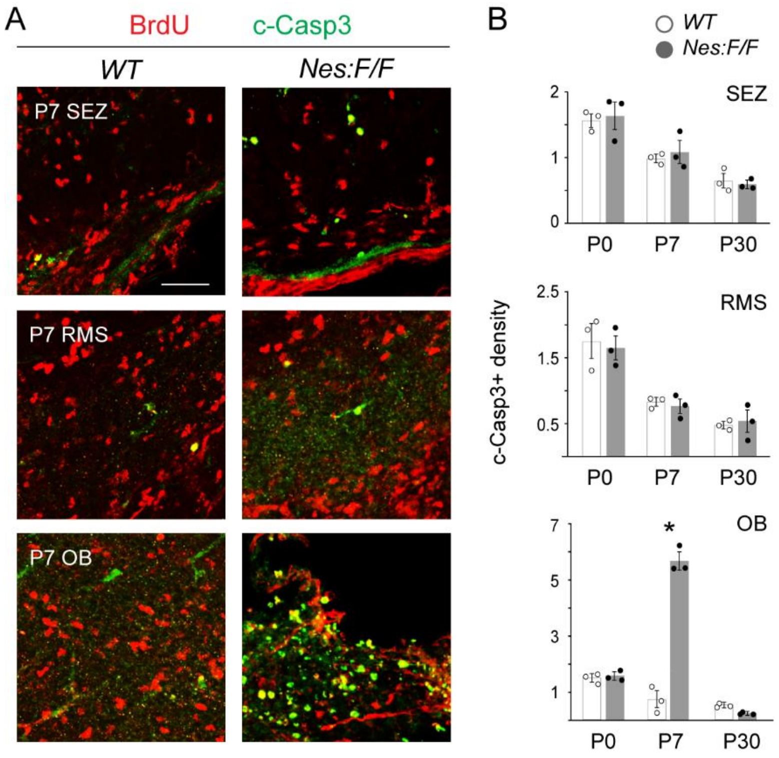


**Figure S5. Analysis of apoptosis in *Nes:F/F* forebrains. Related to Figures 2-3. (A)** Confocal images of BrdU+ and cleaved caspase 3+ (c-Casp3+) cells in the P7 SEZ, RMS, and OB. Scale bars, 50 µm. **(B)** Estimates of c-Casp3+ densities (x $10^4$ / $mm^3$) at P0, P7, and P30 in the SEZ, RMS, and OB of *WT* and control and *Nes:F/F* mice. Data are mean ± s.e.m; individual dots are for each n. Asterisk, Student's t-test, $p < 0.05$.

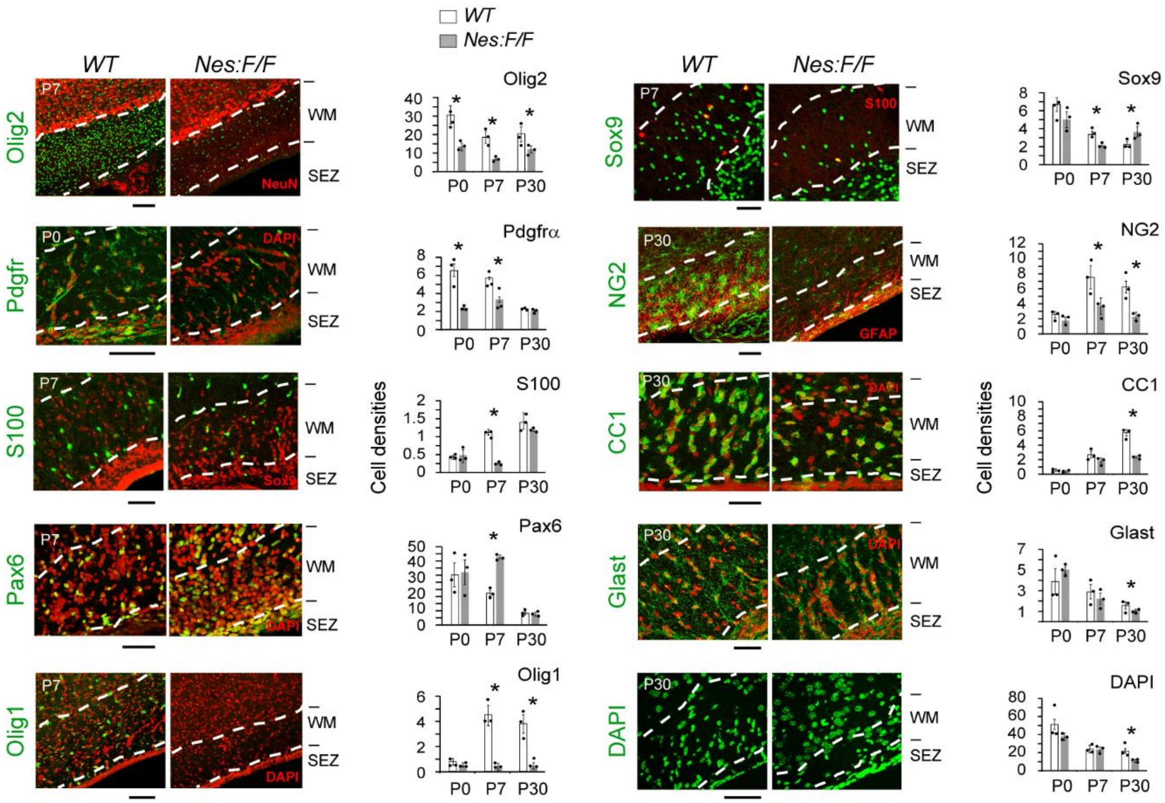


**Figure S6. Marker analysis in the SEZ/WM of the postnatal *Nes:F/F* forebrain. Related to Figures 2-3.** Confocal images of various markers in *WT* and *Nes:F/F* forebrain sections (ages top right) co-stained with DAPI, NeuN, GFAP, Sox9 or S100 as indicated (top right). Dotted lines delineate boundaries of SEZ and WM. Scale bars, 50 µm (apply to matched panels in *Nes:F/F*). Bar charts are density estimates of each marker in the SEZ/WM at P0, P7 and P30 (x $10^4$ / $mm^3$; mean ± s.e.m; individual dots are values for each n). Asterisks, Student's t-test, $p < 0.05$.

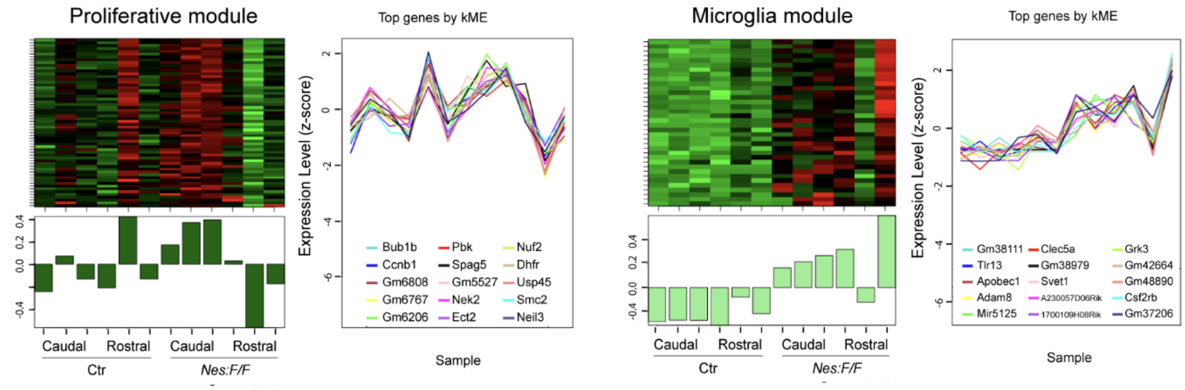


**Figure S7. RNA-seq analysis reveals altered Proliferation and Microglial modules in P5 *Nes:F/F* cortices. Related to Figure 3.** Gene coexpression modules enriched for proliferative and microglia markers reveal differential expression in rostral vs. caudal samples from control (Ctr) and *Nes:F/F* mice at P5. The heatmaps indicate the relative expression levels of the module genes across all samples.  The module eigengene (first principal component) below summarizes its expression pattern. Right plots, z-scored expression levels of the top 15 genes correlated with each module eigengene.

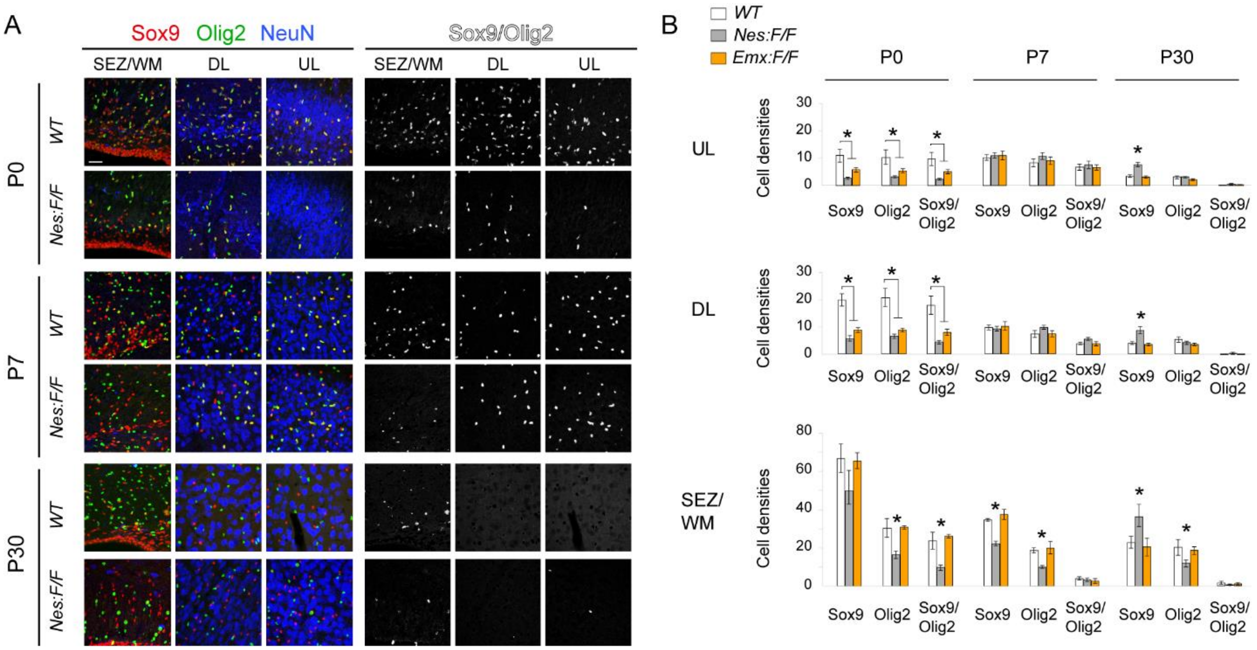


**Figure S8. Co-expression of Sox9 and Olig2+ in dorsolateral cerebral cortices of control and *Nes:F/F* mice. Related to Figure 3. (A)** Confocal images of dorsolateral somatosensory and motor cortices from P0, P7, and P30 *WT* and *Nes:F/F* mice immunostained for Sox9 in combination with Olig2. Bottom gray scale panels are double-labeled nuclei thresholded from the corresponding color panels. Scale bar, 50 µm. **(B)** Cell density estimates ($x10^4$ /$mm^3$) of Olig2+ and Sox9+ singly labeled cells, as well as S100+/ Sox9+ double labeled cells in the cortical layers (subependymal zone, SEZ; white matter, WM; deep layers, DL; superficial or upper layers, UL) at ages and genotypes as indicated. Data are mean ± s.e.m of densities, n=3 animals; Asterisks, $p < 0.05$, Student's t-test.

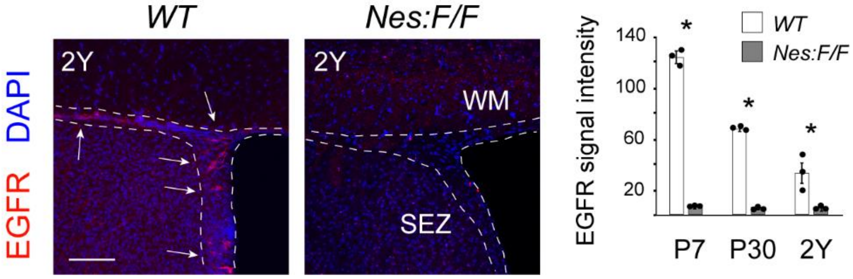


**Figure S9. Persistent loss of EGFR immunoreactivity in the *Nes:F/F* SEZ/WM over the life-span of mice. Related to Figures 2-4.** Confocal images of cortical SEZ/WM regions of 2Y *WT* and *Nes:F/F* mice immunostained for EGFR (red) and counterstained with DAPI (blue). Scale bar, 50 µm. Bar chart are data as mean ± s.e.m of the intensity of EGFR immunoreactivity at P7, P30 and 2Y in the SEZ/WM; individual dots represent values for each n. Asterisks, Student's t-test, $p < 0.05$.

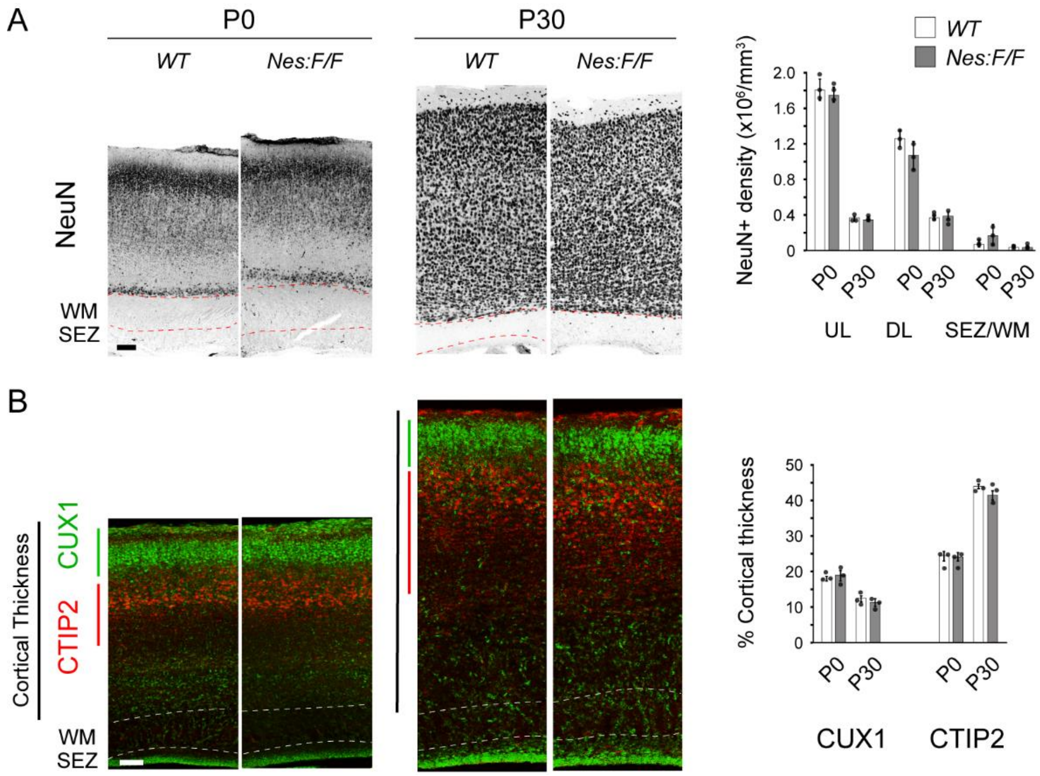


**Figure S10. Neurogenesis and cortical lamination appear intact in *Nes:F/F* cortices. Related to Figures 2-4.** Confocal images of somatosensory and motor dorsolateral cortical layers in *WT* and *Nes:F/F* mice at P0 and P30 immunostained for the neuronal marker NeuN (**A**) as well as CTIP2 and CUX1 (**B**). Scale bars, 100 µm. Bar chart in A are data as mean ± s.e.m of the density of NeuN+ cells. Bar chart in B illustrates percentage of cortical thickness that captures layer-specific distribution of CTIP2+ and CUX1+ cells as indicated. Individual dots in each chart represent values for each n.

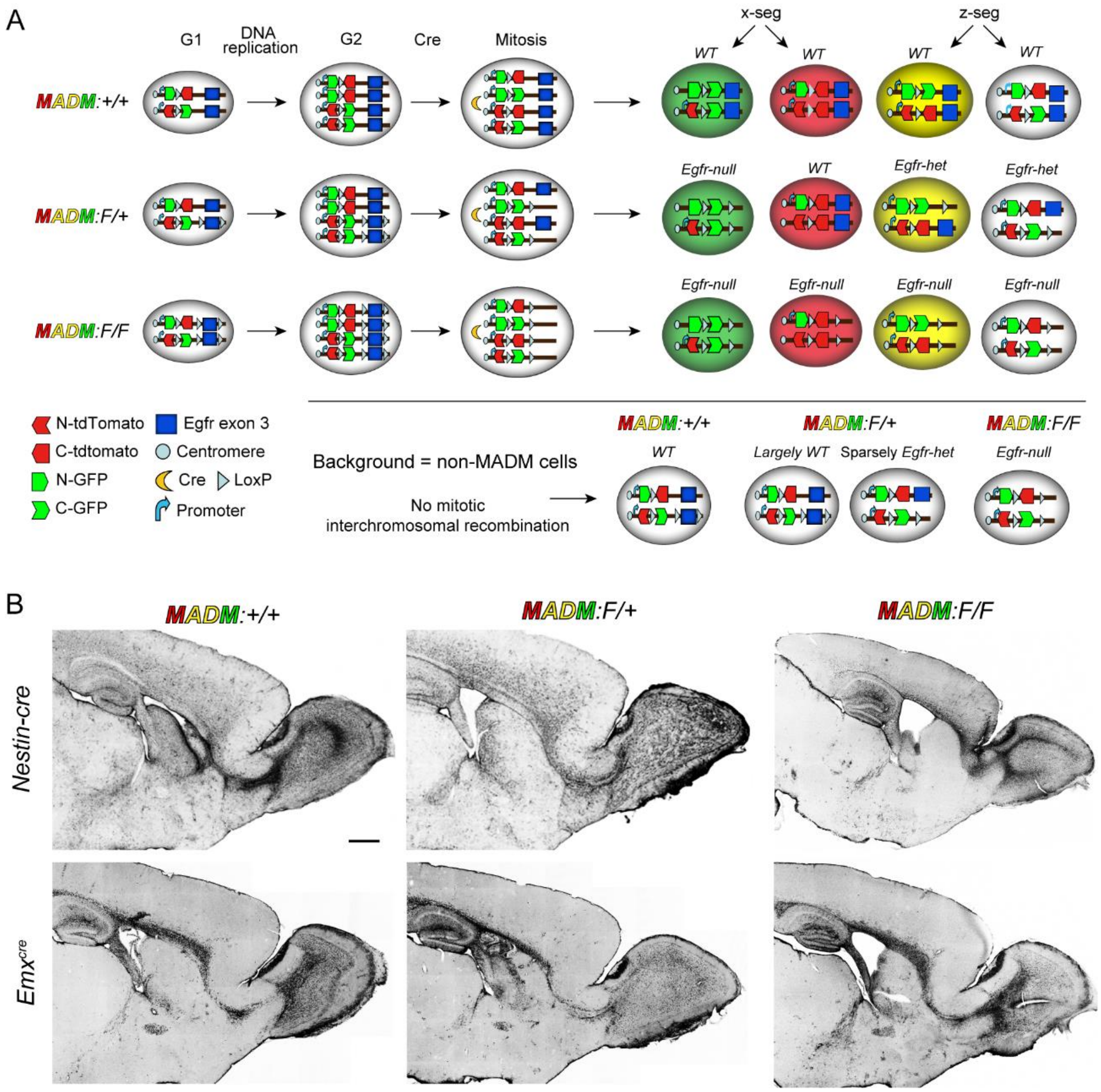


**Figure S11. Mosaic Analysis with Double Markers. Related to Figures 5-7. (A)** Molecular basis of labeling individual progenitor clones with distinct fluorescent reporters (GFP, green and tdTomato, red). Cre mediated interchromosomal mitotic recombination is followed by two segregation patterns (x-seg and z-seg) generating clonal siblings in distinct colors as indicated. In control MADM mice all cells will be wild type (*WT*) and referred to as *MADM:+/+* herein. When a single *floxed* allele for *Egfr* is introduced, green cells lack *Egfr (Egfr-null)* while the red cells are *WT* (*MADM:F/+* mice). A third genotype is obtained when both *floxed* alleles are carried onto the MADM background resulting in both red and green cells with *Egfr-null* genotypes (*MADM:F/F* mice). Background genotypes obtained from the described combinations using cre lines (*Nestin-cre* and *Emx1*$^{cre}$) result in differences in *Egfr* genotype in background cells (non-MADM recombination events) which may exert non-autonomous effects on MADM cells. **(B)** Immunostaining for GFAP to monitor for potential reactive gliosis due to loss of *Egfr* and potential tissue damage. MADM mice at P30 carrying the three genotypes for *Egfr*. Images are sagittal sections. Scale bar, 500 µm.

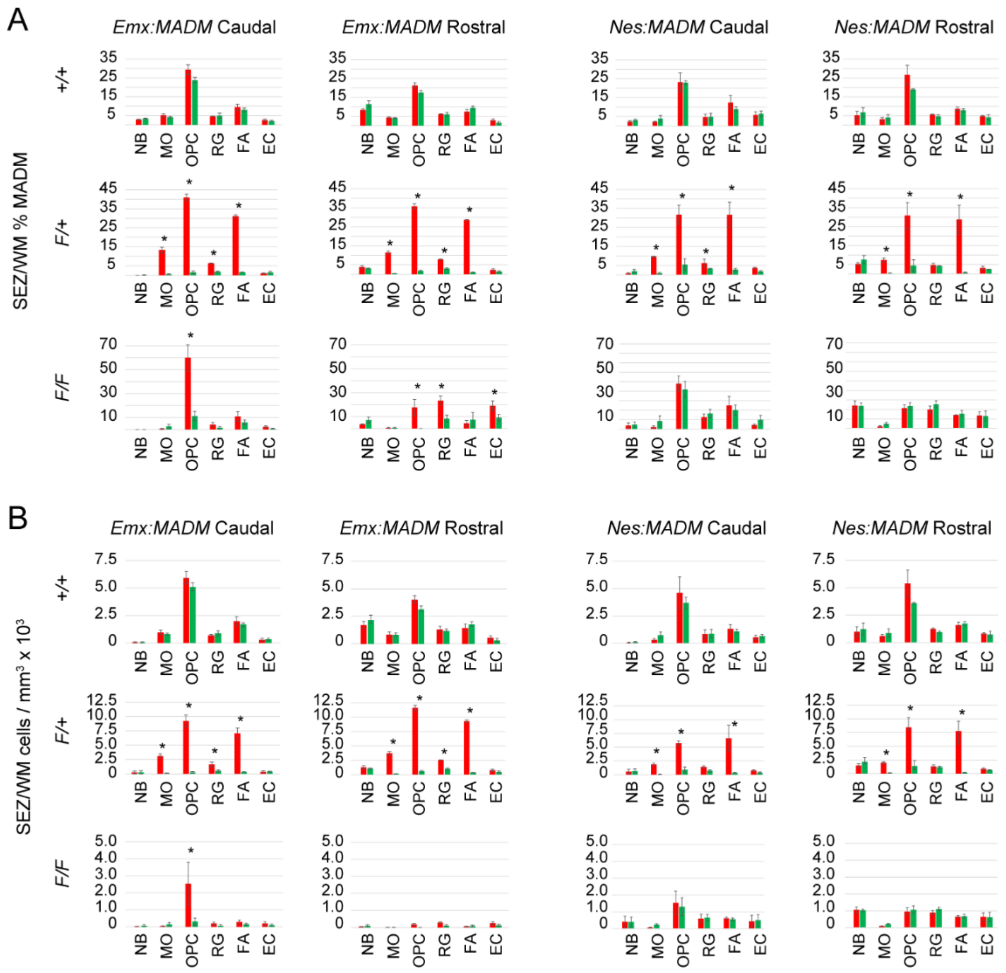


**Figure S12. Analysis of cell types in the SEZ/WM in the MADM forebrain. Related to Figure 6.** (**A**) Percentages and (**B**) densities of red and green MADM cells belonging to each cell type as indicted. Neuroblast (NB), Myelinating oligodendrocytes (MO), Oligodendrocyte Precursor Cell (OPC), Radial Glia (RG), Fibrous Astrocytes (FA), Ependymal Cell (EC). Bar charts are data as mean ± s.e.m; Asterisks, Student's t-test, $p < 0.05$.

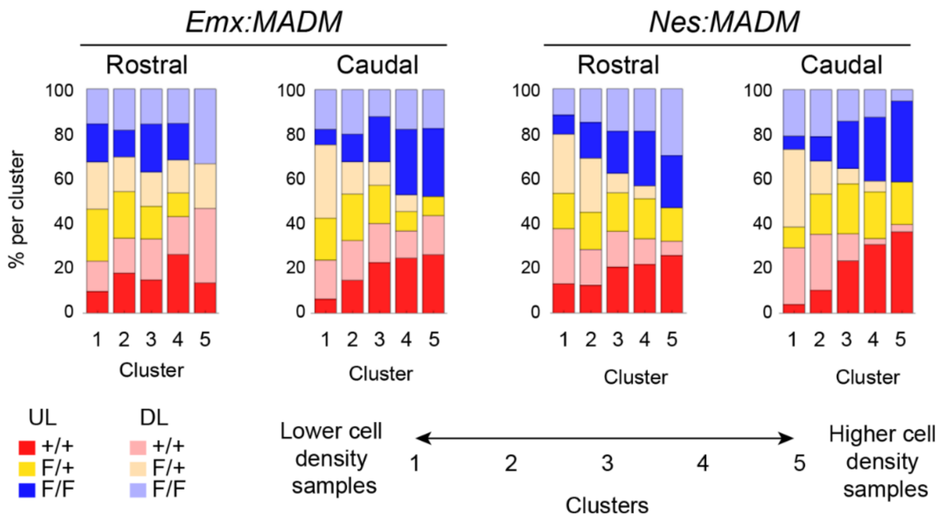


**Figure S13. SOM analysis on neuronal MADM populations. Related to Figure 7.** A self-organizing map unsupervised learning algorithm (SOM) was used to sort neuronal MADM densities into linear arrays of five clusters. Clusterings were performed on neuronal populations in upper (UL) and deep (DL) layers of the cortex as indicated (tdTomato and GFP MADM populations of neurons were pooled in *Emx:MADM* and *Nes:MADM* cortices).

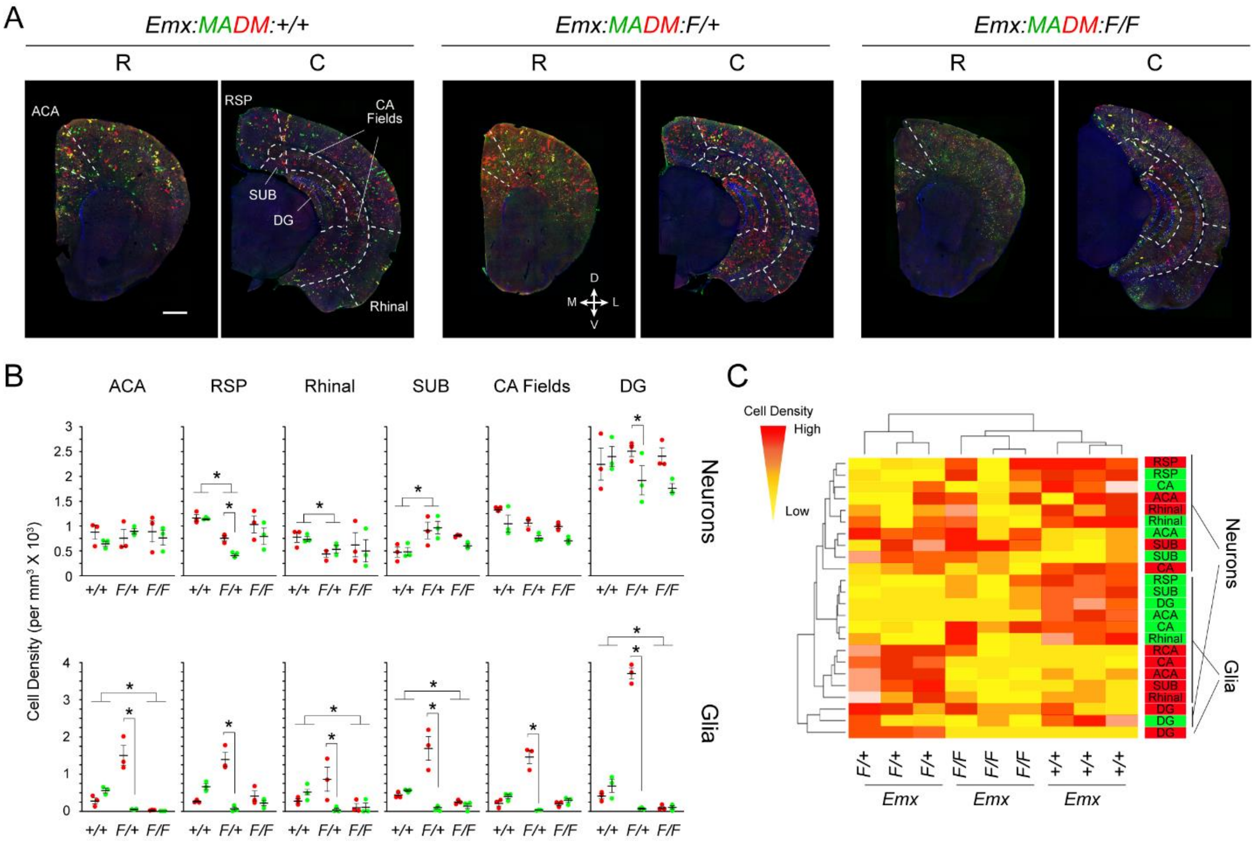


**Figure S14. *Emx:MADM* analysis in paleocortical areas and hippocampal regions interconnected with each other. Related to Figures 5-6. (A)** Representative micrographs through rostral (R) and caudal (C) coronal forebrain sections from P30 mice with the indicated genotypes. Dotted lines demarcate areas where quantifications were conducted: ACA, anterior cingulate area; RSP, retroplenial cortex; Rhinal, entorhinal/perirhinal/ectorhinal cortices; SUB, subiculum; CA Fields, Cornu Ammonis fields; DG, dentate gyrus. Scale bar, 500 µm. **(B)** Cell density measurements of MADM labeled neurons and glia in regions as indicated (mean values x $10^3$ per $mm^3$/ n=3 mice per genotype). Yellow highlight designates significant differences among genotypes in the same regions at $p<0.05$ by pairwise t-test among genotypes or Student's t-test. **(C)** Hierarchical clustering dendrogram of cell density measurements across MADM genotypes per area, cell type and for MADM colors as indicated in the right column. Cell density measurements were used to generate heatmaps with hierarchical clustering. Red signal correlates to higher relative cell densities.
